# Leukemia-intrinsic determinants of CAR-T response revealed by iterative *in vivo* genome-wide CRISPR screening

**DOI:** 10.1101/2022.02.15.480217

**Authors:** Azucena Ramos, Catherine E. Koch, Yunpeng Liu, Riley D. Hellinger, Taeyoon Kyung, Keene L. Abbott, Julia Fröse, Daniel Goulet, Khloe S. Gordon, Rebecca C Larson, John G. Doench, Aviv Regev, Matthew G. Vander Heiden, Marcela V. Maus, Michael E. Birnbaum, Michael T. Hemann

**Affiliations:** Koch Institute for Integrative Cancer Research, Massachusetts Institute of Technology, Cambridge, MA, USA; Department of Biology, Massachusetts Institute of Technology, Cambridge, MA, USA; Department of Biological Engineering, Massachusetts Institute of Technology, Cambridge, MA, USA; Ludwig Center for Cancer Research at MIT, Boston, MA, USA; Broad Institute of Harvard and Massachusetts Institute of Technology, Cambridge, Massachusetts 02142, USA; Klarman Cell Observatory, Broad Institute of MIT and Harvard, Cambridge, MA, USA; Howard Hughes Medical Institute, Chevy Chase, MD, USA; Cellular Immunotherapy Program, Cancer Center, Department of Medicine, Massachusetts General Hospital, Boston, MA, USA; Immunology Program, Harvard Medical School, Boston, MA, USA; Ragon Institute of MIT, MGH, and Harvard, Cambridge, MA, USA

## Abstract

CAR-T therapy is a promising new treatment modality for B-cell malignancies. However, the majority of patients inevitably go on to experience disease relapse through largely unknown means. To investigate leukemia-intrinsic CAR-T resistance mechanisms, we performed genome-wide CRISPR-Cas9 loss-of-function screens in an immunocompetent murine model of B-cell acute lymphoblastic leukemia (B-ALL) utilizing a novel, modular guide RNA library. We identified IFNγ/JAK/STAT signaling and components of antigen processing and presentation pathway as key mediators of resistance to CAR-T therapy *in vivo*, but not *in vitro*. Transcriptional characterization of this model demonstrated an upregulation of these pathways in CAR-T treated relapsed tumors, and examination of data from CAR-T treated patients with B-ALL revealed an association between poor outcomes and increased expression of JAK/STAT/MHC-I in leukemia cells. Overall, our data identify an unexpected mechanism of resistance to CAR-T therapy in which tumor cell interaction with CAR-T cells *in vivo* induces expression of an adaptive T-cell resistance program in tumor cells.

## Introduction

Immunotherapies have emerged as crucial components of cancer treatment, rapidly becoming the standard of care for a broad range of malignancies.^1^ One of the most promising immunotherapy agents today is the adoptive cell transfer (ACT) of autologous T lymphocytes engineered to express chimeric antigen receptors (CARs).^2, 3^ Functionally, CARs redirect the cytotoxicity of immune cells towards a patient’s tumor. Groundbreaking trials in relapsed B cell malignancies demonstrated extraordinary initial efficacy, with upwards of 90% of patients experiencing complete responses (CR).^4–8^ However, despite impressive results, long term follow up data from CAR-T treated patients suggest that relapse is likely to be a significant and ongoing problem in this treatment modality. Recent studies have shown that while the median overall survival after CAR-T therapy is on the order of 12 to 20 months, upwards of 60% of patients will experience disease recurrence ^5, 8–13^ Furthermore, up to 20% of patients with B-ALL never achieve remission and applications in other CD19^+^ B-cell malignancies such as chronic lymphocytic leukemia (CLL) and diffuse large B-cell lymphoma (DLBCL) have significantly lower CR rates.^5, 8, 11, 14^ Thus, gaining a more complete understanding of the factors governing response in CAR-T therapy may ultimately lead to improvements in CAR-T cell design or combination therapies that maintain remission and improve patient outcomes.

To date, a limited number of CAR-T resistance mechanisms have been described.^15^ In some settings, treatment failure has been associated with loss of the infused CAR-T cell product, particularly in patients who never achieve remission.^16, 17^ This has led to the hypothesis that intrinsic CAR-T cell dysfunction is a central determinant of treatment failure due to factors such as the quality of harvested T cells or variations in production and manufacturing. Alternatively, tumor cell intrinsic alterations have also been associated with resistance, with target antigen loss arguably garnering the most attention.^18, 19^ For anti-CD19 CAR-T therapy, various mechanisms of CD19 loss have been reported including mutations in the *CD19* locus, alternative splicing of CD19 mRNA, and lineage switching.^15^ In this setting, as many as 1 in 4 patients relapse with CD19^-^ disease.^20, 21^ Interestingly, patients can also relapse with CD19^+^ disease. However, decidedly less is known about the underlying determinants of relapse in these cases. Presently, it is unclear if intrinsic CAR-T dysfunction is the main driver of resistance in these patients, or whether pre-existing or acquired tumor cell adaptations can function to facilitate or incite CAR-T dysfunction, independent of CD19 status. In fact, recent reports provide support for the latter. For example, Singh and colleagues showed that impaired death receptor signaling in tumor cells led to CAR-T therapy resistance via the induction of CAR-T cell dysfunction.^22^ Gene expression analysis of patient tumor samples prior to CAR-T treatment demonstrated that expression of death receptor pathway genes differed between patients who ultimately achieved CR and those who did not. This pre-treatment tumor expression profile was then able to successfully predict outcomes in an independent patient cohort suggesting that in some cases, CAR-T therapy resistance may arise from pre-existing tumor cell alterations. Overall, mechanisms of CAR-T resistance and relapse beyond intrinsic CAR-T dysfunction or tumor target antigen loss are poorly understood and yet, likely contribute to cases of treatment failure. Ultimately, gaining a better understanding of these additional resistance mechanisms, particularly those driven by tumor intrinsic alterations, could lead to significantly improved therapeutic efficacy via novel combination strategies and better molecular characterization of CAR-T cell-susceptible tumors.

To systematically investigate mechanisms of leukemia-intrinsic resistance to CAR-T therapy in an unbiased manner, we performed parallel whole-genome *in vitro* and *in vivo* CRISPR/Cas9-mediated loss-of-function (LOF) screens in a transplantable, immunocompetent mouse model of *BCR-ABL^+^* B-cell acute lymphoblastic leukemia (B-ALL).^23, 24^ Here, we utilized an iterative screening approach with a first-of-its-kind modular single guide RNA (sgRNA) library and subsequent validation library. Importantly, this pipeline represents a novel approach to CRISPR/Cas9 screening, particularly for *in vivo* settings. Analysis of *in vivo* screening data and gene expression data from CAR-T treated relapsed B-ALL cells identified IFNγ/JAK/STAT signaling and components of antigen processing and presentation pathways as key mediators of CAR-T resistance. Examination of published patient data provided additional support for these findings. Further characterization of our screening and transcriptional data implicated Qa-1^b^ (encoded by *H2-T23*), the murine homolog of Human leukocyte antigen E (HLA-E), in promoting tumor cell resistance to CAR-T therapy. In fact, we found that addition of blocking antibodies preventing interaction of Qa-1^b^ with its inhibitory receptor significantly extend survival when combined with CAR-T therapy. Overall, our data suggest that *in vivo,* CAR-T and tumor cell interactions influenced by interferon gamma (IFNγ) incite leukemia-intrinsic alterations that ultimately fuel resistance and relapse to CAR-T therapy. These findings suggest an “adaptive” resistance mechanism to cell-based immunotherapy and point to new possibilities for enhancing CAR-T cell efficacy without modifying existing CAR-T cell products.

## Results

### A fully immunocompetent mouse model of BCR-ABL1^+^ B-ALL enables parallel *in vivo* and *in vitro* genome-wide screens for CAR-T resistance

Using an established mouse model of *BCR-ABL1*^+^ B-ALL with a high engraftment rate in immunocompetent, syngeneic recipient mice, we engineered Cas9-expressing cells (Cas9^+^) (Supplementary Figure 1a) with high cutting efficiencies (Supplementary Figure 1b) for use in unbiased genome-wide screens.^23–27^ To determine if an *in vivo* screen for CAR-T resistance using immunocompetent mice was tractable, we examined *in vivo* growth in both wildtype (WT) and Cas9^+^ cells reasoning that if Cas9 expression induced any immunogenic barrier it would manifest as delayed growth kinetics over time. A luciferase^+^ Cas9^+^ clone (20.12), growth matched to its WT parental line *in vitro*, was transplanted into non-irradiated immunocompetent male C57BL/6J (B6) mice. Compared to the WT parental line, no significant growth delays in Cas9^+^ cells were detected *in vivo* in any hematopoietic organ assayed (Supplementary Figure 1c). Parallel experiments were also performed in non-irradiated male B6 mice and immunocompromised NOD-SCID/IL2Rg^−/−^ (NSG) mice. If Cas9 was immunogenic in recipient mice, immunosuppressed NSG mice transplanted with Cas9^+^ cells would have succumbed to disease faster than immunocompetent B6 mice transplanted with the same cells. However, no differences in disease latency were observed in repeated experiments (Supplementary Figure 1d).

We then examined the ability of murine CAR-T cells to suppress tumor cell growth *in vivo*. Clinically, patients treated with CAR-T therapy first undergo lymphodepletion with cyclophosphamide or irradiation. Thus, we subjected recipient mice to an irradiation-based lymphodepletion protocol prior to tumor transplantation and subsequent CAR-T treatment. ^28–30^ To calibrate the dose of CAR-T cells, we performed a series of *in vivo* dosing experiments utilizing CD28-based 2^nd^ generation murine CARs targeting murine CD19 (mCD19). Previous groups utilizing a similar construct successfully suppressed disease with CAR-T dose ranges in the millions of cells (5×10^6^ to 2×10^7^) per animal.^28, 29, 31^ In our model, doses of 7×10^6^ to 1×10^7^ CAR-T cells per animal administered two days after the transplantation of 0.6×10^6^ B-ALL cells achieved significant dose-dependent life extension (Figure 1a). To simultaneously monitor disease suppression in real time, we used leukemia cells engineered to express firefly luciferase.^26^ Bioluminescence imaging completed at multiple time points after ACT demonstrated that anti-mCD19 CAR-T cells could significantly suppress disease over time (Figure 1b-f).

**Figure 1.**
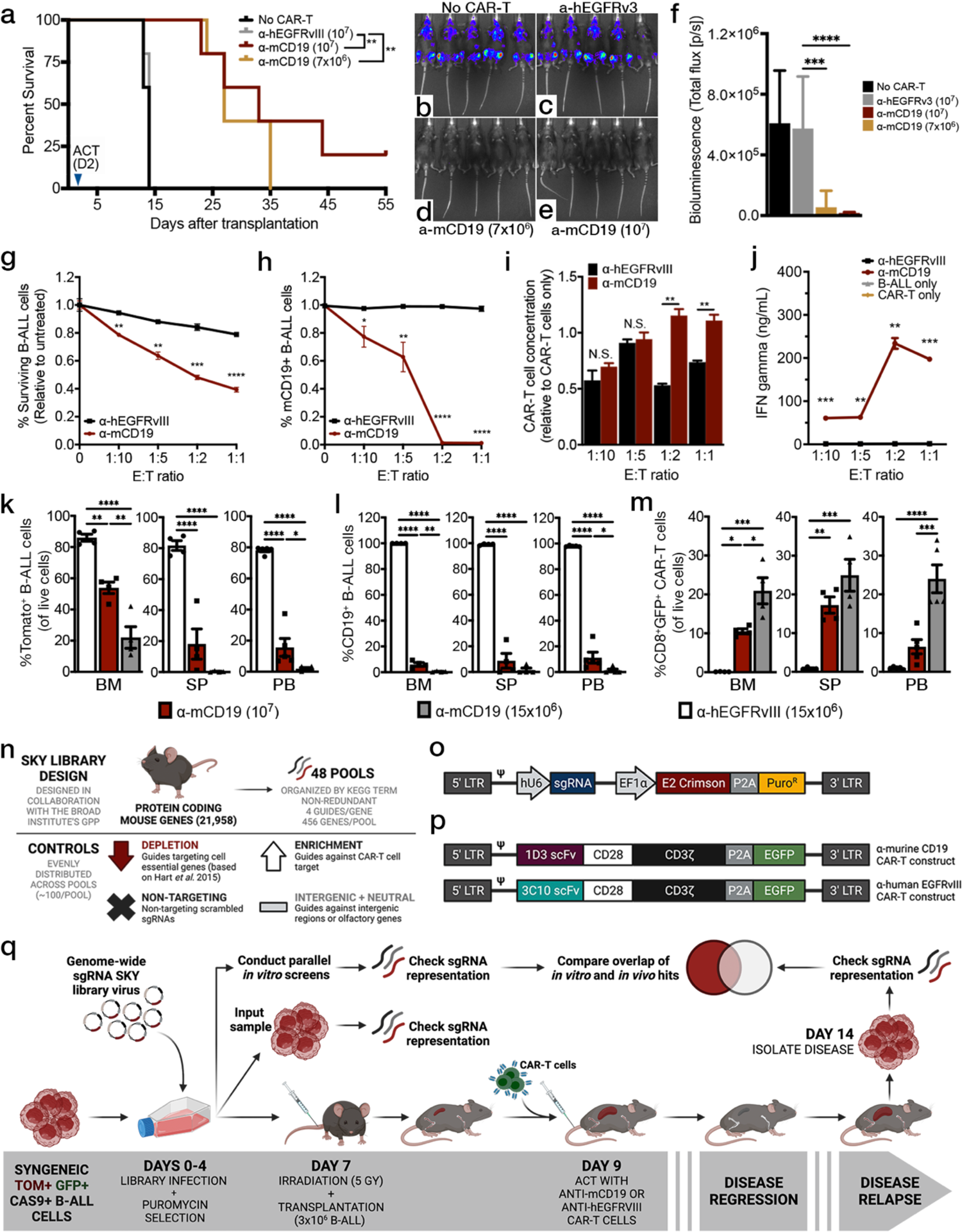
A fully immunocompetent mouse model of BCR-ABL1^+^ B-ALL enables parallel *in vivo* and *in vitro* genome-wide screens for CAR-T resistance. (a) Survival curves and analysis of irradiated B6 mice inoculated with syngeneic B-ALL cells and treated with indicated CAR-T cell type. Mice treated with CAR-T cells targeting murine CD19 (mCD19) showed significant life extension in a dose-dependent manner, as compared to those treated with no CAR-T cells or control CAR-T cells against human EGFRvIII (hEGFRvIII or hEGFRv3). Bioluminescence imaging four days after adoptive cell transfer (ACT) showed significant disease burden in (b) mice not treated with CAR-T cells and (c) mice treated with control CAR-T cells. Conversely, mice treated with mCD19 CAR-T cells at a dose of either (d) 7×10^6^ or (e) 10^7^ show (f) significant disease suppression. (g) *In vitro* cytotoxicity assays show significant depletion of tumor cells along with (h) massive target epitope loss when B-ALL cells are treated with anti-mCD19 CAR-T cells at increasing effector to target cell (E:T) ratios, as assayed via flow cytometry. (i) Concomitantly, CAR-T cells expand and (j) release IFNγ when co-cultured with mCD19+ B-ALL cells. (k-m) Experiments to determine the appropriate dose of CAR-T cells for either the bone marrow (BM) or spleen (SP) using flow cytometry analysis. Peripheral blood (PB) was also assessed. Mice sacrificed four days after ACT showed an 80% reduction in bone marrow disease at a dose of 1.5×10^7^ anti-mCD19 CAR-T cells, while identical disease suppression in the spleen and blood was accomplished with 10^7^ CAR-T cells. (l) Massive target epitope loss and (m) significant CAR-T cell persistence was also observed in all of the organs harvested from mice treated with anti-mCD19 CAR-T cells. (n) Schematic showing the overall design and (o) lentiviral backbone used to create the SKY library. (p) Retroviral vectors encoding the 1D3 single chain variable fragment (scFv) targeting mCD19 (top) and the 3C10 scFv targeting hEGFRvIII (bottom). (q) Diagram of screening layout. Data are mean ± s.e.m.; n = 4–6 mice per group. All experiments were repeated at least twice with representative data shown. Significance for survival experiments was determined using log-rank tests. For all other experiments, significance is determined using unpaired two-sided student’s t-tests with Bonferroni correction for multiple comparisons or using one-way ANOVA with Turkey’s correction for multiple comparisons when more than two groups were compared. *P<0.05; **P < 0.01; ***P < 0.001; ****P < 0.001.

To ensure CAR-T functionality, we conducted concurrent matched *in vitro* cytotoxicity assays for each independent experiment and measured CAR-T expansion and IFNγ (a cytokine released by T cells in proinflammatory conditions) in the resulting culture supernatant. Interestingly, B-ALL cell numbers could be significantly reduced but never fully eliminated *in vitro* even at very high effector to target (E:T) ratios (Figure 1g). CAR-T cells also induced a rapid and dramatic loss of the mCD19 target epitope on the surface of B-ALL cells (Figure 1h), a phenotype that our results suggest is target antigen independent and not a unique feature of our murine B-ALL model (Supplementary Figure 1e). Importantly, while antigen loss was a primary phenotype in our B-ALL cells, our cytotoxicity assays still resulted in activated CAR-T cells that expanded (Figure 1i) and released significant IFNγ after being co-cultured with cells expressing their target antigen (Figure 1j).

Our group has previously shown that this particular mouse model of B-ALL is highly amenable to *in vivo* LOF screens.^25, 26^ Thus, we next sought to determine the appropriate CAR-T cell dose to use for *in vivo* screening.^25, 26^ Irradiated B6 mice were transplanted with B-ALL cells (clone 20.12), treated with varying CAR-T cell doses, and monitored using bioluminescence imaging. Animals were sacrificed at peak disease suppression which occurred on or before day three after CAR-T treatment (Supplementary Figure 1f). Total disease burden, target tumor antigen expression, and CAR-T expansion were assayed in various hematopoietic organs. We aimed for 80-90% disease suppression, as this significant but incomplete level of tumor cell reduction would allow for identification of alterations that could sensitize tumor cells to therapy but would not completely eradicate tumor cells. For the bone marrow compartment, this was accomplished using 1.5×10^7^ CAR-T cells, while splenic and peripheral blood compartments only required a dose of 1×10^7^ CAR-T cells (Figure 1k). Relapsed disease harvested from CAR-T treated mice showed a striking dose-dependent loss of mCD19 surface expression along with significant CAR-T engraftment in all organs examined (Figure 1l-m), a consistent phenotype observed across all *in vivo* experiments. Given that we also observed significant antigen loss and concomitant CAR-T cell persistence in relapsed animals (Supplementary Figure 1g), we reasoned that this phenotype was due to ongoing CAR-T surveillance. Indeed, when mCD19-leukemia cells were harvested from relapsed mice and cultured *in vitro*, mCD19 expression was restored as CAR-T cells were depleted from culture (Supplementary figure 1h-i). Notably, clinical data has demonstrated that as many as half of all patients relapse with antigen negative disease. Thus, our system may model the antigen loss and leukemia relapse seen following CAR-T therapy in a large proportion of patients.^5,9,32^

### Unbiased CRISPR/Cas9-mediated screen identifies genes and pathways involved in *in vivo* resistance to anti-CD19 CAR-T therapy

Having established a mouse model of anti-mCD19 CAR-T treatment response, we next sought to develop a CRISPR/Cas9 library that would enable genome-wide *in vivo* screens across a broad range of tumor models with varying engraftment rates. We generated a novel, genome-wide pooled sgRNA library cloned into an optimized lentiviral backbone containing a crimson fluorophore and puromycin selection marker (Figure 1n-p). The library targets each protein coding gene (total of 21,958 genes) in the mouse genome with 4 different sgRNAs per gene. Genes are organized by KEGG term and evenly distributed into 48 individual, non-redundant sub-pools (∼456 genes/pool). All sgRNAs targeting a given gene are present in the same sub-pool and there are control sgRNAs evenly distributed across all pools. This unique feature allows for sub-pools to be used as stand-alone screening libraries that can also be combined into larger sgRNA pools, depending on the model system. Given the high *in vivo* engraftment rate of our B-ALL model, we were able to pool our 48 sub-pools into groups of eight, limiting our entire genome-wide *in vivo* and *in vitro* screens to six separate screens completed over two experiments (Figure 1q).

To query factors responsible for CAR-T resistance, we subjected library-infected Cas9^+^ B-ALL cells (20.12) grown *in vitro* or *in vivo* to either anti-mCD19 CAR-T therapy or control CAR-T cells targeting human EGFRvIII (hEGFRvIII) which is not expressed in mice or on our B-ALL model. The *in vivo* arm of our screen followed the layout of our optimized dose finding experiments. We observed that relapsed CAR-T treated mice harbored both CD19-positive and -negative disease, as well as persistent CAR-T cell populations in both the spleen and bone marrow (Supplementary Figure 2a and b, respectively). Concurrent with this *in vivo* screen, we performed parallel *in vitro* screens at two effector to target (E:T) cell ratios (Figure 1q) completed on the same timeline as *in vivo* screens. Tumor cells were treated with CAR-T therapy on day 9 and maintained in culture until isolation on day 14. To assay CAR-T functionality during the primary screen, IFNγ release assays were performed after 24 hours of co-culture. In all cases, CAR-T cells released significantly more IFNγ when exposed to their target antigen and no significant differences between experiments could be detected (Supplementary Figure 2c). On day 14, live, sgRNA-bearing B-ALL cells were isolated from animal organs or cell culture using fluorescence activated cell sorting (FACS) (Supplementary Figure 2d). We then used high-throughput sequencing to quantify sgRNA representation in B-ALL cells harvested from each experimental arm. Importantly, we were able to confirm that we achieved extensive coverage of every sgRNA library pool screened (Supplementary Figure 2e-f). To confirm that our screen was able to effectively interrogate gene function while remaining unaffected by possible artifacts from Cas9-mediated DNA breakage, we quantified the enrichment and depletion of sgRNAs in the control treatment group compared to pre-screen input samples. Indeed, sgRNAs targeting known essential genes significantly depleted in the control treatment groups, whereas those targeting intergenic regions as well as non-targeting sgRNAs did not (Supplementary Figure 2g).

As expected, samples treated with anti-mCD19 CAR-T therapy demonstrated patterns of sgRNA enrichment and depletion that were largely distinct from control treated groups (Supplementary Figure 2h). To assess the genetic dependencies involved in the response of B-ALL cells to anti-mCD19 CAR-T treatment, we compared the anti-mCD19 CAR-T and control treatment arms of the screens across each of the six pools (Figure 2a). These results were then aggregated to generate a genome-wide perturbation landscape of CAR-T treatment response (Figure 2b-c). Within such a landscape, depleting sgRNAs target genes that putatively mediate resistance to CAR-T therapy, whereas those that enrich target genes that likely sensitize leukemic cells to CAR-T therapy. Two central features support the biological relevance of our findings: first, sgRNAs targeting the *Cd19* locus are the top overall enrichers (Figure 2b-d) and second, the magnitude of enrichment and depletion appears to be positively associated with the anti-mCD19 CAR-T cell dose administered (i.e., the selective pressure experienced by target cells) (Figure 2b, left and middle panels and Figure 2c). Interestingly, a comparison of top hits between the *in vivo* and *in vitro* arms of our screen (Figure 2e) showed little overlap, suggesting that an *in vitro* co-culture screening modality is incapable of capturing key *in vivo* mechanisms of CAR-T treatment resistance. Furthermore, pairwise comparisons of hits between each condition confirm this finding showing that while screen results obtained in the same context are highly correlated (Supplementary Figure 3a, *in vivo* vs. *in vivo* and *in vitro* vs. *in vitro* highlighted in red and blue boxes, respectively), *in vitro* results are poorly correlated to those generated *in vivo* (Supplementary Figure 3a, *in vitro* vs. *in vivo* highlighted in the gold box). In fact, certain sgRNAs exhibit opposite behavior in *in vivo* versus *in vitro* settings. For example, top enrichers in the *in vitro* arms of the screen including sgRNAs targeting the IFNγ signaling pathway members Janus kinase 2 (*Jak*2), signal transducer and activator of transcription proteins 1 (*Stat1*), and interferon gamma receptor 1 (*Ifngr1*) deplete in the *in vivo* arms of the screen (Figure 2b-d).

**Figure 2.**
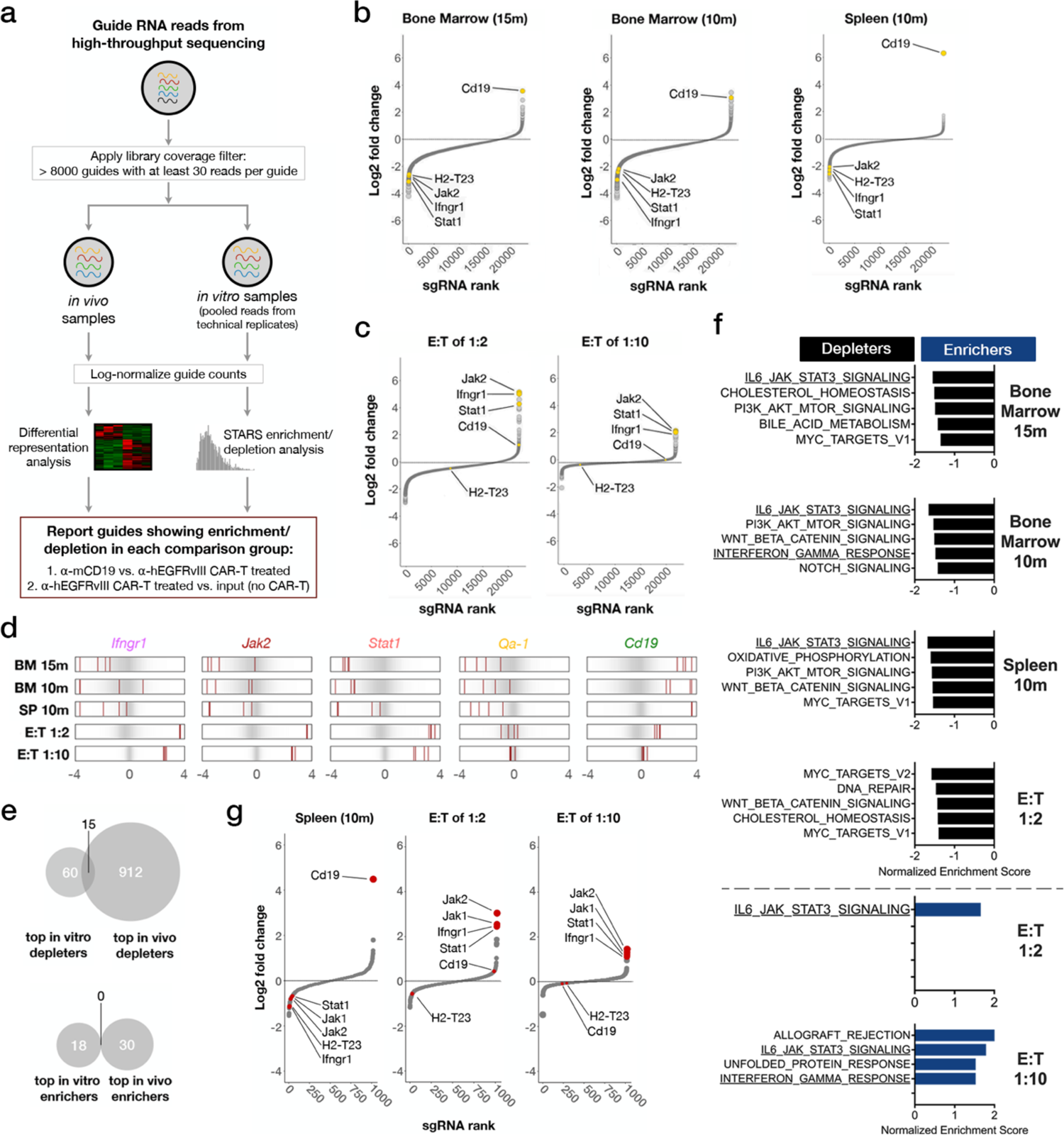
*In vivo* genome-wide primary and subsequent validation CRISPR-Cas9-mediated knockout screens identify IFNγ/Jak/Stat signaling and antigen presentation pathways involved in resistance to CAR-T therapy. (a) Schematic of the data analysis and screen hit discovery workflow. (b-c) Waterfall plots of log-fold changes of the representation of sgRNAs against different genes in anti-mCD19 CAR-T cell treated animals (b) or cells (c) compared to anti-hEGFRvIII CAR-T cell treated (control) animals (b) or cells (c) at indicated doses (15m or 1.5×10^7^ CAR-T cells, 10m or 1×10^7^ CAR-T cells, effector to target (E:T) ratios of 1:2, or 1:10) during primary screens. For *in vivo* screens, the organ from which guide-bearing B-ALL cells were collected is also indicated. Genes are ranked by the average log-fold changes of all sgRNAs against each gene, and point sizes are proportional to the magnitude (absolute value) of log-fold changes. Waterfall plots display the relative significance of each hit (vertical axis, showing log2(fold change) from two-sided student’s t-test with correction for multiple comparisons or STARS analysis) versus guide ranks. (d) Relative enrichment/depletion of individual guides against top depleters *Ifngr1*, *Jak2*, *Stat1,* and *Qa-1* are shown in each arm of the screen. Guide RNAs against *Cd19* are also shown as an indicator of CAR-T treatment pressure. (e) Venn diagrams showing overlap of top hits from the primary *in vitro* and *in vivo* arms of the screen. Top hits are selected based on two criteria: 1) the strongest average signals (based on absolute log-fold change) and 2) most stable hits across all mice in the group, as scored by coefficient of variation (CV). The final list is a union of genes scoring highest by either criterion over each of the 6 individual pools. (f) Bar plots of GSEA analysis of top depleting (black) or enriching (blue) hits in each arm of the primary screen. Plots show up to the top 5 Hallmark pathways with enrichment FDR value smaller than 0.05 enriched Hallmark pathways. (g) Waterfall plots from the validation screen showing the representation of sgRNAs against different genes in anti-mCD19 CAR-T cell treated animals (spleen) or cells compared to anti-hEGFRvIII CAR-T cell treated (control) animals (spleen) or cells. Waterfall plots display the relative significance of each hit (vertical axis, showing log2(fold change) from STARS analysis) versus guide ranks. Guides against *Cd19* and against top *in vivo* depleters (*Ifngr1*, *Jak1, Jak2*, *Stat1,* and *Qa-1*) are also shown. All screens were performed once, with multiple biological (*in vivo*) or technical (*in vitro*) replicates, as indicated in Supplemental Figure 2e-f and Supplemental Figure 3d.

We next sought to determine whether pathways independent of target molecule (CD19) expression were involved in resistance to CAR-T therapy. Gene set enrichment analysis (GSEA) revealed that several signaling and metabolic processes within the MSigDB hallmark gene sets, such as the JAK/STAT/IL-6 signaling axis, the PI3K/AKT pathway, and oxidative phosphorylation (OXPHOS) were enriched among genes mediating resistance to CAR-T therapy *in vivo* (Figure 2f, depleters, above dotted line). In contrast, some of these pathways were enriched among genes that potentially sensitize leukemic cells to CAR-T therapy *in vitro* (Figure 2f, below dotted line). Additionally, KEGG pathway enrichment analysis (Supplementary Figure 3b) showed that top *in vivo* depleters were enriched for processes such as tumor necrosis factor (TNF) signaling, immune checkpoint signaling, and immune response against infectious diseases. Interestingly, the interferon gamma receptor 1 (IFNGR1) pathway, along with components of JAK/STAT/IL-6 signaling and the murine histocompatibility 2, T region locus 23 (*H2-T23*, also known as *Qa-1^b^*) are components of multiple enriched pathways. Accordingly, guides against these genes are among the top *in vivo*-specific depleters (Figure 2b, d).

Our initial genome-wide pool-based screening approach made direct comparisons between sgRNAs across pools difficult. To enable a more definitive ranking of all sgRNAs exhibiting significant biological effects, we selected top hits from each pool and created a validation library containing six newly generated, unique sgRNAs per gene, targeting a total of 933 genes. We also included 385 non-targeting (scrambled) guides, 100 intergenic cutting control guides, and 135 guides with consistently neutral behavior in our initial screens. In order to minimize any differences due to CAR-T production over the two initial experiments, validation library genes were selected from the top 5% of genes in each pool (divided evenly between enrichers and depleters) based on two criteria: 1) sgRNAs exhibiting the largest fold differences in representation relative to input averaged among all animals and 2) the most stably enriched or depleted sgRNAs across all mice in the group, as scored by the coefficient of variation (CV). The final hit list is a union of genes scoring by either criterion over each of the six individual pools. A validation screen completed in a manner identical to the primary screens was then performed. As in our primary screens, guides targeting known essential genes significantly depleted in the control treatment groups relative to all other guides (Supplementary Figure 3c) and *in vivo* samples treated with anti-mCD19 CAR-T cells exhibited patterns of sgRNA enrichment and depletion distinct from the control group (Supplementary Figure 3d). As expected, sgRNAs targeting the *Cd19* locus were among the top overall enrichers *in vivo* (Figure 2g, left panel). Guides targeting the IFNγ/JAK/STAT signaling pathway were once again amongst the top depleting sgRNAs *in vivo* and amongst the top enrichers *in vitro*, suggesting that these genes may be central negative regulators of the cellular response to CAR-T therapy *in vivo* (Figure 2g).

### The IFNγ pathway promotes resistance to CAR-T therapy *in vivo*

To further validate our findings, particularly hits in the IFNγ/JAK/STAT pathway, we conducted parallel *in vivo* and *in vitro* competition experiments (Figure 3a-c and Supplementary Figure 4a-b). Here, we used a new, independently generated Cas9-expressing cell line (RH62) from our parental B-ALL model with high cutting efficiency and matched *in vitro* growth kinetics to our screened clone (Supplementary Figure 4c-d). We focused on genes whose loss sensitized tumors to CAR-T therapy *in vivo*, as they represent novel drug targets that could potentiate the effects of CAR-T therapy. Using a fluorescence-based competition assay comparing the relative growth of mixtures of isogenic knock out or control B-ALL cells *in vitro* and *in vivo*, we found that the proportion of cells null for *Ifngr1*, *Jak2*, or *Stat1* significantly depleted in both the bone marrow and spleens of mice treated with anti-mCD19 CAR-T therapy (Figure 3b, c showing log-scale depletion). Notably, no growth differences were observed in mice transplanted with identical cell mixtures treated with control CAR-T cells, indicating that this phenotype was not driven by fitness defects imparted on cells lacking these genes.

**Figure 3.**
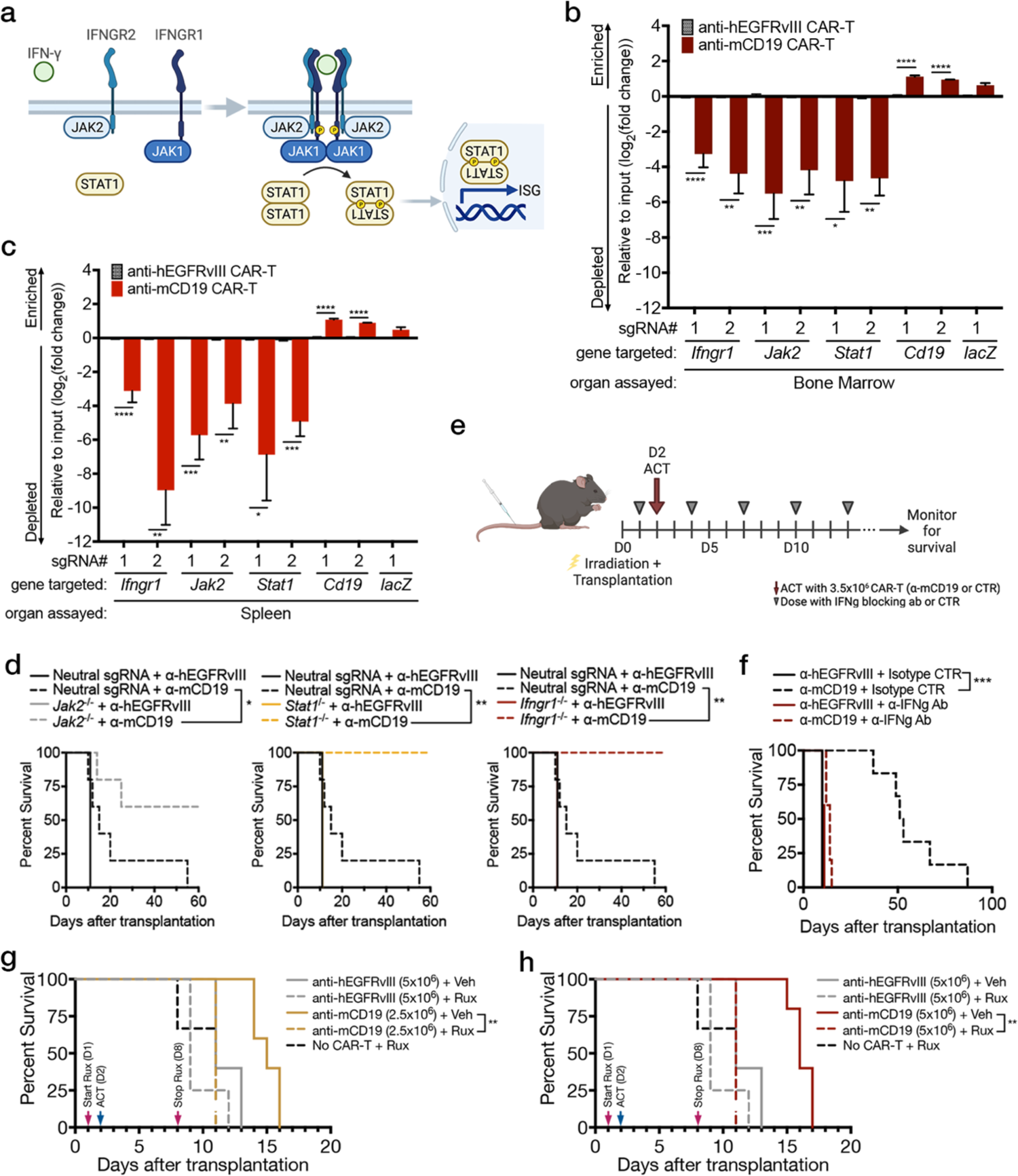
Loss of components of the IFNγ/JAK/STAT pathway sensitizes tumors to CAR-T therapy *in vivo*. (a) Schematic showing the IFNγ signaling pathway. (b-c) *In vivo* competitive assays demonstrate specific log-fold depletion of Cas9^+^ RH62 B-ALL cells lacking components of the IFNγ/JAK/STAT pathway and enrichment of B-ALL cells lacking mCD19 after treatment with anti-mCD19 CAR-T cells in both (b) the bone marrow and (c) the spleen. (d) Immunocompetent mice transplanted with B-ALL cells deficient in indicated components of the IFNγ/JAK/STAT pathway survive significantly longer than control mice. (e) Layout and treatment schedule for anti-IFNγ blocking antibody experiment. (f) Blocking IFNγ signaling concurrently with CAR-T therapy eliminates the antitumor capabilities of CAR-T cells. Concurrent treatment of Ruxolitinib, a JAK1/2 inhibitor, with either 2.5×10^6^ (g) or 5×10^6^ (h) CAR-T cells also eliminates any benefit of CAR-T therapy. *In vivo* competition assays were repeated three times. Survival experiments were completed twice. Pharmacologic studies using anti IFNγ blocking antibody or JAK inhibitor were completed once each. Significance for survival experiments was determined using log-rank tests. For all other experiments, significance is determined using unpaired two-sided student’s t-tests with Bonferroni correction for multiple comparisons. Data are mean ± s.e.m.; n = 5-8 mice per group. *P<0.05; **P < 0.01; ***P < 0.001; ****P < 0.001.

Additionally, observed results were not related to Cas9 immunogenicity, as *Ifngr1^-/-^*cells also significantly depleted in transgenic Cas9 mice transplanted with identical isogenic cell mixtures and treated with CAR-T therapy derived from transgenic Cas9 mice (Supplementary Figure 4e). Interestingly, parallel *in vitro* experiments again demonstrated the opposite result, with *Ifngr1*, *Jak2*, or *Stat1* knock out cells significantly enriching in the context of anti-mCD19 CAR-T therapy (Supplementary Figure 4a). Ultimately, these data confirm our screen findings that loss of *Ifngr1*, *Jak2*, or *Stat1* sensitizes B-ALL cells to CAR-T therapy *in vivo* while promoting resistance *in vitro*.

To determine whether our findings were specific to leukemia cells, we examined the effect of *Ifngr1* deficiency on CAR-T therapy response in a murine model of glioblastoma. Here, we transduced the extracellular domain of human CD19 (hCD19) into GL261 cells and performed intra-cranial injections into syngeneic recipient mice. As in the B-ALL competition experiment, we mixed fluorescently labeled GL261 cells deficient for *Ifngr1* with isogenic WT control cells and tracked the relative behavior of each cell type following injection of anti-hCD19 CAR-T cells. Similar to the B-ALL setting, we observed significant depletion of *Ifngr1* knock out cells suggesting that our results are not limited to one tumor type (Supplementary Figure 4f).

Next, to further validate our *in vivo* B-ALL results, we generated pure, clonal populations of leukemic cells deficient in *Ifngr1*, *Jak2*, or *Stat1* (Supplementary Figure 5a). Consistent with our previous data, mice transplanted with these cells demonstrated increased sensitivity to anti-mCD19 CAR-T therapy resulting in significantly increased survival and complete lack of tumor relapse or detectable disease during the indicated time period following treatment (Figure 3d and Supplementary Figure 5b). Thus, tumor cells can directly co-opt IFNγ signaling *in vivo* to resist CAR-T cell killing. Conversely, *in vitro* CAR-T efficacy appears to be at least partially dependent on the ability of tumor cells to sense IFNγ. Together, these data highlight the significant difference between *in vitro* and *in vivo* contexts and demonstrate the importance of investigating tumor cell response to CAR-T therapy in a physiologically relevant setting. These results also underscore a central role for the IFNγ/JAK/STAT pathway in promoting resistance to CAR-T therapy in B-ALL.

### Global targeting of JAK/STAT signaling does not enhance CAR-T therapy

Having established the IFNγ/JAK/STAT pathway as a key mediator of response to CAR-T therapy, we next explored whether global blocking of IFNγ *in vivo* could potentiate the effects of CAR-T therapy. To this end, we administered blocking antibodies targeting IFNγ the day before ACT, and every three days thereafter (Figure 3e). Rather than enhancing antitumor effects, blocking IFNγ in the context of anti-mCD19 CAR-T treatment abrogated the anti-tumor effects of CAR-T cells. Mice treated with anti-mCD19 CAR-T and anti-IFNγ antibody succumbed to disease at the same time as mice treated with control CAR-T cells and an isotype control antibody (Figure 3f and Supplementary Figure 5c). Identical results were obtained when animals were co-treated with ruxolitinib (a JAK1/2 inhibitor) and CAR-T therapy in a similar experiment (Figure 3g, h and Supplementary Figure 5d-j). These data are consistent with results from recent studies which show that treating CAR-T cells with JAK/STAT inhibitors or knocking out *Ifng* in CAR-T cells significantly impairs their ability to eliminate tumor cells *in vitro* and *in vivo*, respectively.^31, 33^ Thus, IFNγ is indispensable for sustaining CAR-T cell cytotoxicity in an immunocompetent *in vivo* context, resulting in the lack of a therapeutic window for targeting this cytokine directly to sensitize tumor cells to CAR-T cell killing.

### *H2-T23* is an *in vivo*-specific mediator of resistance to CAR-T therapy

To begin to explore how IFNγ/JAK/STAT signaling promotes resistance to CAR-T therapy, we examined screen hits downstream of this pathway with known immunoinhibitory functions. One of the most promising genes was the non-classical class I major histocompatibility complex (MHC-I) gene *H2-T23* which encodes Qa-1^b^, the murine homolog of human leukocyte antigen E (HLA-E).^34, 35^ Surface expression of HLA-E has been shown to be induced by IFNγ signaling.^36, 37^ In addition, HLA-E is the only known ligand of the CD94/NKG2A receptor which is expressed on the surface of natural killer (NK) cells and CD8^+^ T-cells.^38–42^ Binding of NKG2A/CD94 to its ligand transmits a signal that inhibits the effector functions of NK and CD8^+^ T cells (Figure 4a).^36, 40–43^ Recently, several groups have reported significant enhancement in the antitumor effects of immune checkpoint blockade (ICB) and cancer vaccines when combined with blockade of the HLA-E-NKG2A axis, and early clinical trials have demonstrated encouraging results in a variety of cancers, including hematologic malignancies.^35, 36, 43, 44^ Notably, sgRNAs targeting *H2-T23* showed potent *in vivo* depletion in our primary genome-wide screen and were among the top depleters in our validation screen.

**Figure 4.**
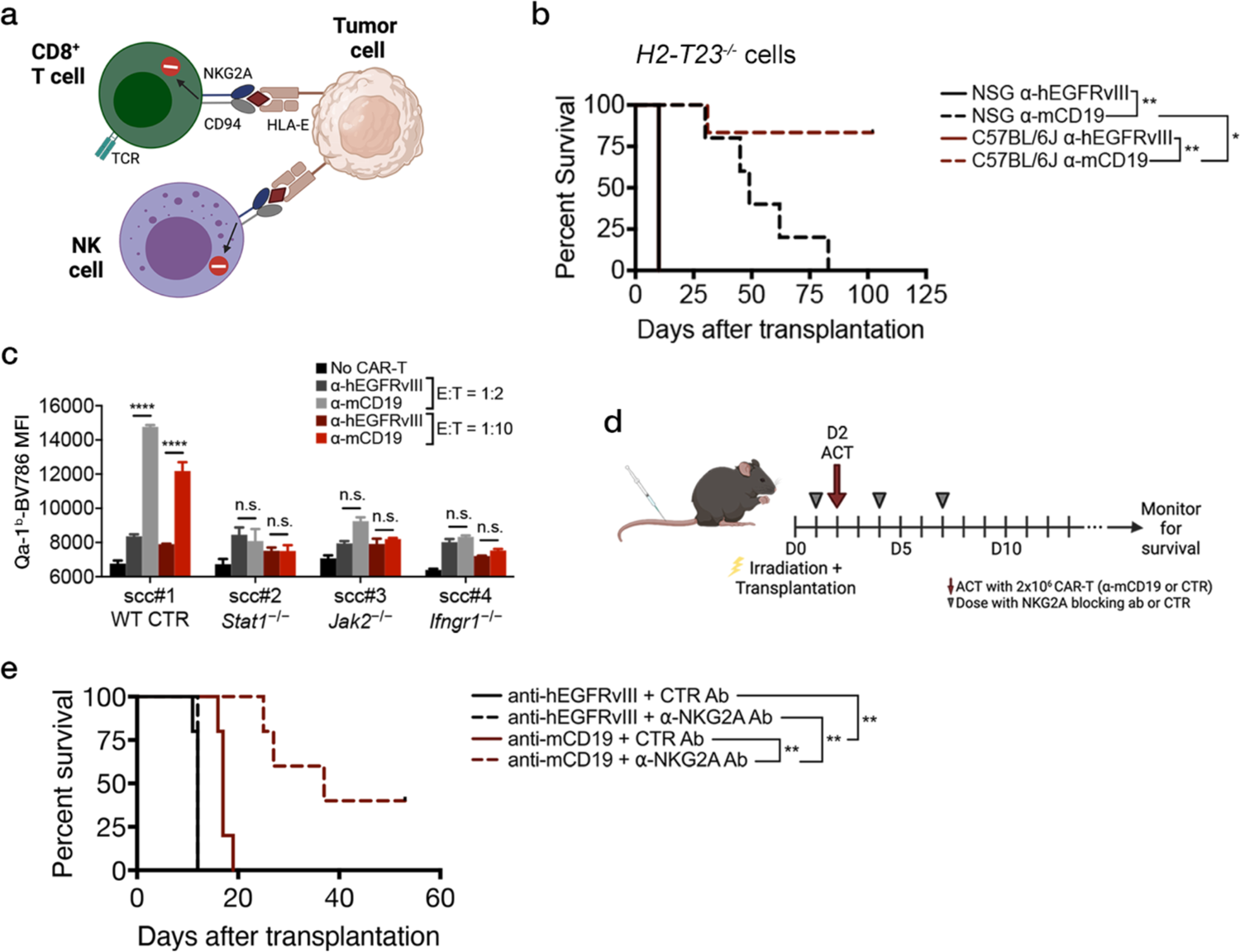
Loss of Qa-1^b^, a component of the MHC-I pathway, or pharmacologic blockade of NKG2A, the only known receptor of Qa-1^b^, sensitizes B-ALL cells to CAR-T therapy *in vivo*. (a) Schematic showing the known effect of the HLA-E/NKG2A/CD94 axis in human NK and CD8^+^ T-cells. (b) Immunocompetent (B6) mice transplanted with Qa-1^b^-deficient B-ALL cells show increased survival as compared to immunocompromised (NOD/SCID/IL2Rγ or NSG) mice transplanted with the same *H2-T23^−/−^* B-ALL cells and treated with anti-mCD19 CAR-T cells. (c) Mean fluorescence intensity (MFI) of Qa-1^b^-Brilliant Violet 786 (BV786) after *in vitro* CAR-T treatment of single cell clones (scc) deficient in indicated components of the IFNγ/JAK/STAT pathway at two different effector to target (E:T) cell ratios, as assayed using flow cytometry. Wildtype control single cell clones were also generated and assayed at the same time. Representative data for WT control clones is shown in (c). (d) Layout and treatment schedule for anti-NKG2A blocking antibody experiment. (e) Blocking the NKG2A/Qa-1^b^ axis with a murine version of the anti-NKG2A antibody Monalizumab, concurrently with CAR-T therapy potentiates the antitumor capabilities of CAR-T cells, leading to increased survival *in vivo*. The survival experiment in (b) and *in vitro* CAR-T treatment of *H2-T23^−/−^* cells in (c) were completed at least twice and were paired each time. Representative data is shown. The pharmacologic experiment in (e) was completed once. Significance for survival experiments was determined using log-rank tests. For all other experiments, significance is determined using unpaired two-sided student’s t-tests with Bonferroni correction for multiple comparisons. Data are mean ± s.e.m.; n = 5 mice per group. *P<0.05; **P < 0.01; ***P < 0.001; ****P < 0.001.

To further examine the consequence of Qa-1^b^ loss on tumor cell response to CAR-T therapy, we generated pure populations of Qa-1^b^-null B-ALL lines by sorting for bulk sgRNA-bearing cells that could no longer be induced to express Qa-1^b^ (Supplementary Figure 6a-c).^43^ Consistent with our screen results, Qa-1^b^ knockout tumors were sensitized to CAR-T therapy *in vivo* leading to significant life extension, including an absence of tumor relapse in fully immunocompetent animals (Figure 4b). Notably, immunodeficient NSG recipient mice bearing Qa-1^b^-deficient tumors showed extended survival in response to CAR-T therapy but, unlike B6 recipients, eventually relapsed suggesting that full CAR-T efficacy may require additional cellular components of the adaptive immune system. When tested *in vitro* using fluorescence-based competition assays as before, no changes in the proportion of Qa-1^b^ knockout cells could be detected in any treatment group, as demonstrated in our primary screen (Supplementary Figure 6a). Finally, to investigate whether cells deficient in components of the IFNγ/JAK/STAT pathway were also functionally null for Qa-1^b^, we performed *in vitro* cytotoxicity experiments. While control cells were fully capable of inducing dose-dependent expression of Qa-1^b^ upon exposure to anti-mCD19 CAR-T cells, isogenic cell lines deficient in *Ifngr1*, *Jak2*, or *Stat1* were completely blunted in their ability to express this molecule at all E:T ratios tested (Figure 4c) or in repeated experiments using recombinant IFNγ in lieu of CAR-T therapy (Supplementary Figure 6b-d).

To determine whether interruption of HLA-E-NKG2A inhibitory signaling might have therapeutic potential in the context of CAR-T therapy, we introduced an antibody targeting NKG2A into leukemia-bearing mice concurrently with the introduction of CAR-T cells (Figure 4d-e and Supplementary Figure 6e). This antibody blocks Qa-1^b^-mediated inhibitory signaling through NKG2A. Recipients of the anti-NKG2A antibody showed significantly longer leukemia-free survival relative to control treated animals, including a number of mice failing to relapse over the course of the experiment.

While NKG2A is present on subsets of CD8^+^ T-cells, it is best characterized as an inhibitory NK cell receptor. Given that *H2-T23* deficient tumor cells relapsed after CAR-T treatment in immunodeficient NSG mice lacking NK cells but not in immunocompetent B6 mice possessing NK cells, we hypothesized that NK cells may be contributing to anti-leukemia activity in the context of CAR-T therapy. To examine the role of NK cells in this setting, we depleted NK cells in B6 mice using an antibody targeting NK1.1, an established method for *in vivo* NK cell depletion.^45–47^ NK1.1 is an activating surface receptor expressed on almost all NK cells in mice and its human homolog CD161 is expressed on a majority of human NK cells.^48–50^ Leukemia-bearing mice were treated with anti-mCD19 or control CAR-T therapy and dosed concurrently with NK1.1 or isotype control antibody. Mice were dosed again with NK1.1 or isotype control antibodies seven days later in order to maintain NK cell depletion (Supplemental Figure 4f, g).

No difference in survival was observed between mice receiving control CAR-T treatment and NK1.1 antibody or isotype control antibody suggesting that NK depletion alone has no effect on B-ALL disease progression. Surprisingly, mice receiving anti-mCD19 CAR-T treatment and NK1.1 antibody had significant survival extension compared to mice treated with anti-mCD19 CAR-T therapy and isotype control suggesting that the addition of NK1.1 antibody improves CAR-T antitumor efficacy. One possible explanation for this is activation of other NK1.1 expressing cells, such as Natural Killer T-cells (NK-T cells) or subsets of CD4^+^ and CD8^+^ T-cells (perhaps even some NK1.1^+^ CD8^+^ CAR-T cells).^50^ Various groups have reported populations of NK1.1^+^ cells that are refractory to *in vivo* depletion by NK1.1 antibody and instead become activated.^50, 51^ Alternatively, it is also possible that NK cells negatively impact CAR-T function, leading to improved efficacy when they are depleted. While additional experiments must be performed to obtain a definitive answer, these findings reveal a potential therapeutic strategy for improving CAR-T efficacy through combination with NK1.1 antibody. Additionally, these data suggest that Qa-1^b^ on tumor cells most likely blocks CAR-T efficacy directly via interaction with NKG2A/CD94 present on CAR-T cells rather than on NK cells.

### A IFNγ/JAK/STAT program delineates cells refractory to anti-CD19 CAR-T therapy

To further explore transcriptional programs driving B-ALL CAR-T resistance mechanisms highlighted by our screen and validation experiments, we transcriptionally profiled B-ALL cells after *in vivo* treatment with CAR-T cells. Principal component analysis of transcription profiles using the most variable genes across samples showed a clear separation of animals treated with anti-CD19 CAR-T cells from those treated with control anti-hEGFRvIII CAR-T cells (Figure 5a). Differential expression analysis showed that several genes involved in JAK/STAT signaling and antigen processing and presentation pathways, including *Stat1*, *Irf1*, *Socs1*, and several MHC genes, were highly expressed in samples treated with anti-CD19 CAR-T therapy compared to those treated with control CAR-T therapy (Figure 5b). Interestingly, sgRNAs targeting these genes were also among the top depleters in our *in vivo* screens, suggesting that these genes are part of an expression program that may contribute to therapeutic resistance. Consistently, GSEA of Hallmark pathways highlighted interferon response, allograft rejection, MYC/2F targets, and IL6/JAK/STAT3 signaling as top pathways enriched in samples treated with anti-CD19 CAR-T cells (Figure 2c).

**Figure 5.**
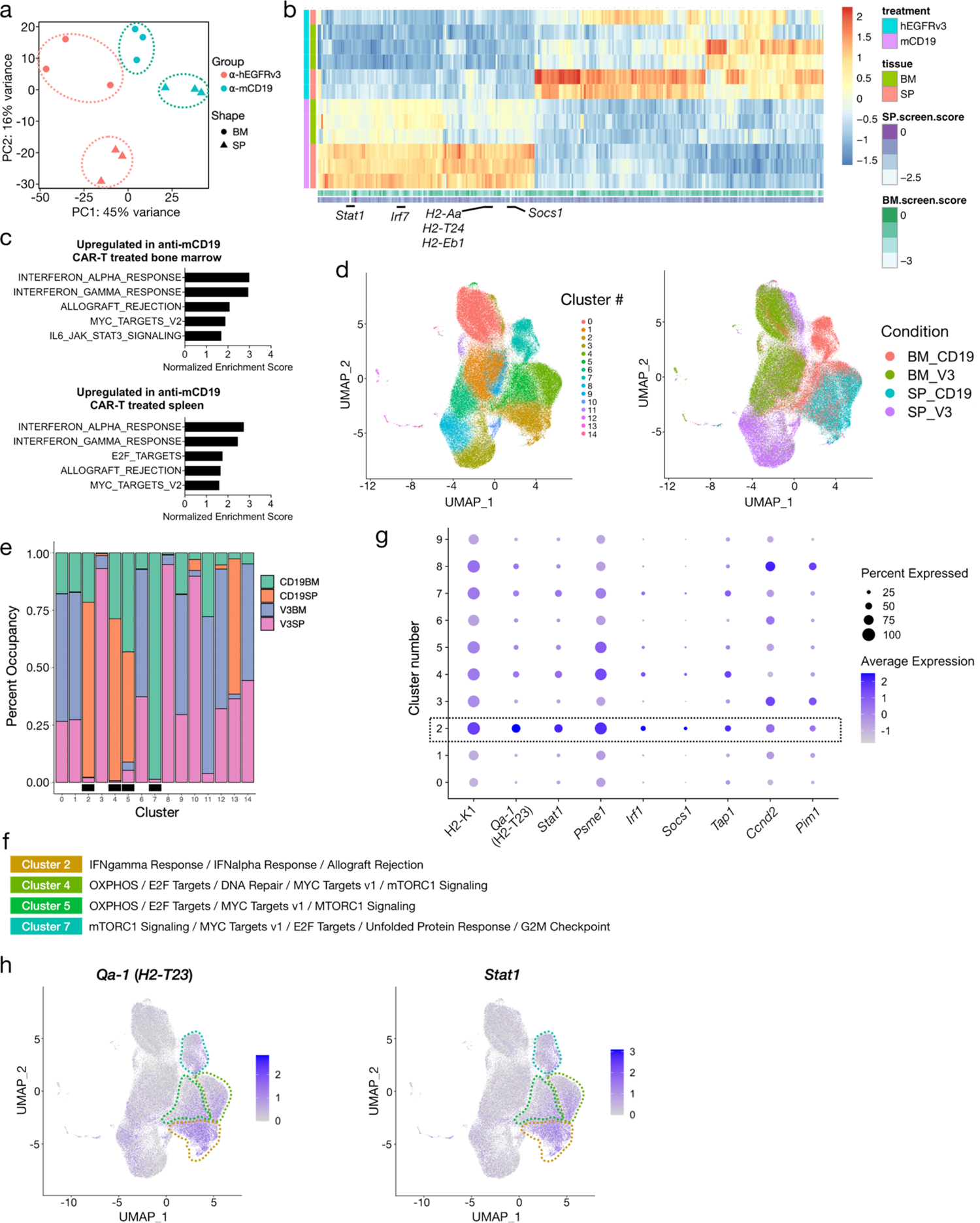
Bulk and single cell gene expression profiling pinpoints specific cell subsets and expression programs as a source of CAR-T cell resistance in B-ALL. (a) PCA plot of bulk RNA-seq profiles of bone marrow (BM) and spleen (SP) samples collected from mice treated with either anti-mCD19 or anti-hEGFRvIII (control) CAR-T cells. (b) Heatmap of genes differentially expressed between anti-mCD19 and anti-hEGFRvIII CAR T cell-treated samples. Scores of genes from the screen are shown next to each gene’s expression profile (column). Top depleters in the bone marrow and spleen samples are labeled. (c) Top Hallmark gene sets enriched among genes overexpressed in mCD19 CAR T cell therapy compared to control treated bone marrow or spleen samples. Bar graphs show normalized enrichment scores for each of the top 5 enriched pathways. Enrichment analysis was performed using GSEA and all pathways shown have FDR < 0.05. (d) 2-dimensional UMAP plots of single cell gene expression profiles collected from mice treated with either anti-mCD19 or anti-hEGFRvIII CAR T cell therapy. Data are collected from n = 3 mice per treatment group. Left panel: cells color-coded by clusters discovered through unsupervised clustering. Right panel: cells color-coded by treatment and tissue groups. (e) Participation of cells from each tissue/treatment group in each cluster. Bar graphs show percentage of cells in each cluster belonging to each tissue/treatment group (anti-mCD19 CAR-T cell treated bone marrow or spleen labeled CD19BM and CD19SP, respectively; anti-hEGFRvIII control CAR-T treated bone marrow or spleen labeled V3BM and V3SP, respectively). Clusters where there is substantial enrichment of cells under anti-mCD19 CAR T therapy are marked with a black box underneath the bar plots. (f) Hallmark gene sets enriched in the 4 treatment-specific clusters as labeled in (e). (g) Dot plot showing relative expression patterns of cluster 2 marker genes which are also among top depleters (log2 fold change < −1.75) in the BM samples of the *in vivo* screen. Dot size is proportional to the percentage of cells in each cluster expressing each gene and dot color indicates average expression of each gene in each cluster. (h) UMAP plots showing expression of *Qa-1^b^* and *Stat1* in single cell samples, with cells color-coded by expression levels. All RNA sequencing experiments (bulk and single cell) were completed once with n = 3 mice per group.

Similar to many other tumor types, B-ALL often displays a heterogeneous response to therapy *in vivo*. To dissect the *in vivo* transcriptional responses to anti-CD19 CAR-T therapy at a finer resolution and pinpoint specific cell populations giving rise to relapse, we performed droplet-based single cell RNA-seq (scRNA-seq) on a total of 124,523 cells harvested from the bone marrow and spleen of mice treated with anti-CD19 or control anti-hEGFRvIII CAR-T cells. Unsupervised clustering revealed 15 distinct cell populations as seen in the 2-dimensional uniform manifold approximation and projection (UMAP) plots in Figure 5d (left panel). Cells from different treatment arms and tissues occupy distinct clusters (Figure 5d, right panel and Supplementary Figure 7a). Interestingly, we did not observe loss of *Cd19* transcript expression in any sample (Extended Data Figure 7b, left panel) indicating that loss of CD19 surface expression in this context is likely the result of post-translational regulation. Remarkably, a few clusters (2, 4, 5, and 7) were substantially enriched for cells from anti-CD19 CAR-T treatment groups while being depleted of cells from the control groups (Figure 5e). Consistent with our bulk RNAseq data, these clusters were marked by elevated expression of genes in the IFNγ signaling and allograft rejection pathways, as well as MYC and E2F target sets (Figure 5f). Importantly, several signature genes in cluster 2 were also among the top depleters in our *in vivo* screens. Figure 5g highlights the magnitude and pervasiveness of the expression of these genes across cell clusters. Additionally, clusters 4, 5, and 7 were enriched for genes involved in OXPHOS and mTORC1 signaling pathways (Figure 5f). Cells in clusters 2 and 4, which contain mostly splenic disease from anti-CD19 CAR-T treated animals, showed widespread elevated expression of *Stat1* and *H2-T23* (Figure 5g, h). Intriguingly, transporter associated with antigen processing (*Tap1*) is also over-expressed in these clusters (Supplementary Figure 7b, right panel). Cluster 7, which is almost entirely comprised of cells from anti-CD19 CAR-T treated bone marrow, is distinguished by a strong G2/M arrest phenotype (Figure 5f and Supplementary Figure 7c) along with elevated expression of *H2-T23* and *Stat1* (Figure 5h). Transcription factor (TF) motif analysis of gene regulatory regions specifically upregulated in clusters 2, 4, 5 and 7 revealed enrichment of motifs critical for TFs involved in interferon response such as ETS family TFs, IRF8, and ELK4, with particularly strong enrichment in clusters 2 and 4 (Supplementary Figure 7d).

Our single-cell and bulk expression data, taken together with results from our *in vivo* screens and validation assays, further support the hypothesis that cross talk between antigen processing and presentation, and interferon gamma signaling plays a crucial role in dictating B-ALL response to anti-CD19 CAR-T therapy. Here, a subset of cells characterized by elevated expression of genes associated with these pathways may be responsible for therapeutic resistance.

### High JAK/STAT signaling correlates with CAR-T resistance in humans

Finally, to examine whether JAK/STAT signaling in tumor cells might also promote resistance to CAR-T therapy in human B cell malignancies, we examined RNAseq data sets generated using pre-treatment bone marrow biopsy samples from patients with B-ALL who received CAR-T therapy.^22^ We generated a “sensitizer signature” from the overlap of our bone marrow depleting hits and reasoned that decreased expression of this signature should correlate with better outcomes in patients since these genes represent novel resistance mediators to CAR-T therapy. In line with our prediction, patients who experienced complete responses (CRs) show significantly less expression of this “sensitizer signature” compared to non-responders (NRs) (Figure 6a). To further examine these patient data, we also generated a JAK/STAT/MHC-I resistance signature by overlapping the top depleters in our screen with the top overexpressed genes in tumor cell cluster 2 from our transcriptional analysis. Strikingly, this resistance signature was correlated with poor outcomes, with NRs demonstrating significantly increased expression of our JAK/STAT/MHC-I gene set compared to CRs (Figure 6b). Notably, no correlation was seen between pre-treatment HLA-E gene expression and clinical outcome (Supplementary Figure 8), likely due to the fact that IFNγ/JAK/STAT signaling induces HLA-E peptide loading and cell surface occupancy rather than alterations in gene expression. These expression data are consistent with recent reports in large B cell lymphoma (LBCL) where high intratumoral interferon signaling is correlated with poor outcomes after CAR-T treatment in patients.^52^ Here, tumor cell expression of an interferon stimulated gene resistance signature (ISG.RS) associated with ICB resistance, is a strong predictor for CAR-T treatment failure in LBCL.^53, 54^ Further examination of this signature in our model revealed that relapsed B-ALL cells with high IFNγ signaling after CAR-T treatment failure were also enriched for expression of the ISG.RS gene set (Figure 6c). Taken together, these data suggest that intratumoral IFNγ/JAK/STAT signaling, along with downstream antigen processing and presentation pathways, may be key determinants of CAR-T response in human B-ALL. Notably, our screens can also nominate novel combination strategies to re-sensitize tumor cells to CAR-T therapy, such as concurrent blockade of the inhibitory receptor NKG2A (Figure 6d).

**Figure 6.**
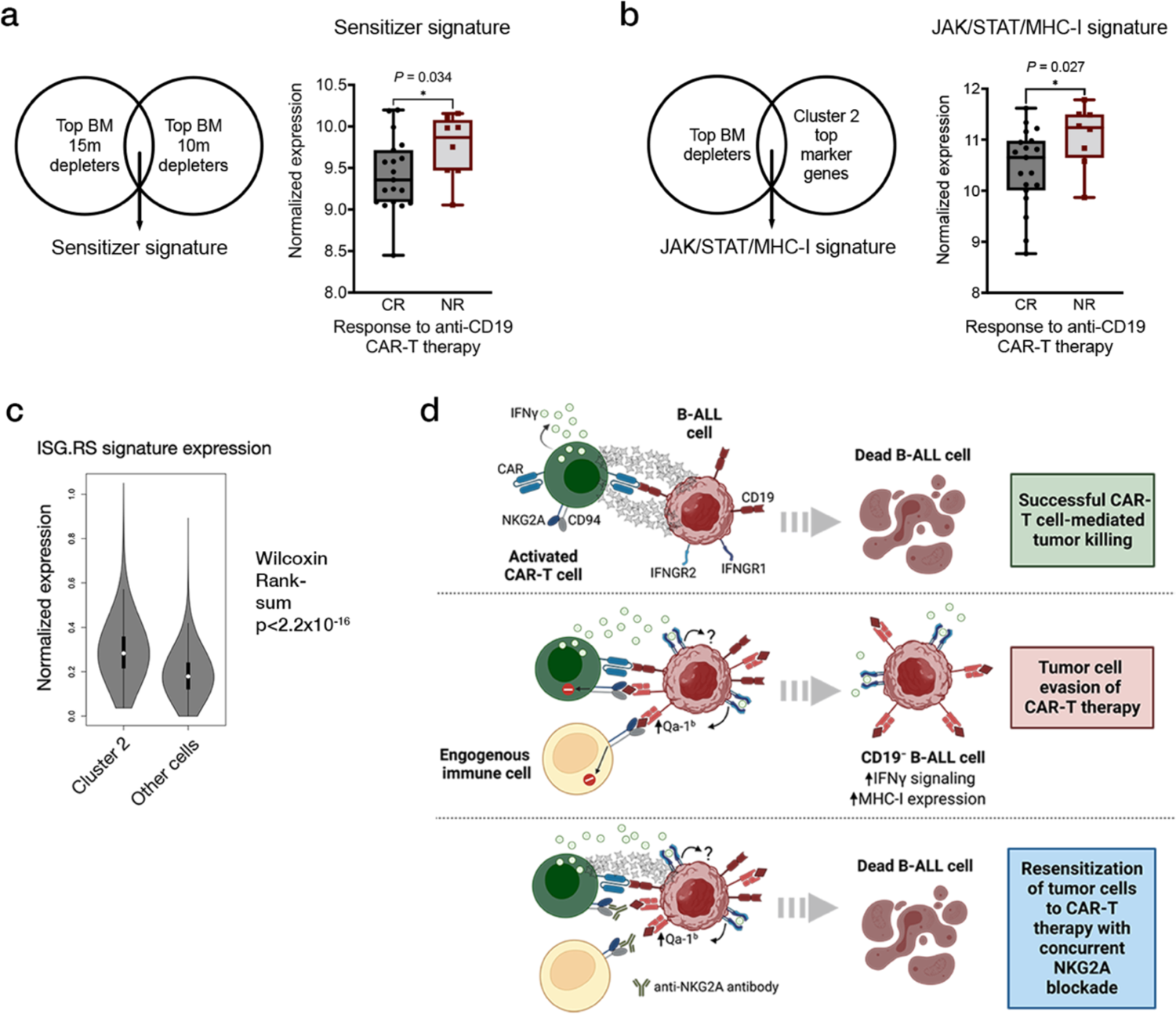
JAK/STAT signaling is a potential therapeutic target in human B-ALL that can be exploited to enhance CAR-T therapy. (a) A gene signature composed of top depleting genes in the bone marrow is expressed at a lower level in complete responders (CR) to anti-mCD19 CAR-T therapy and is thus associated with better outcomes in patients with B-ALL. Non-responders (NR) show increased expression of these genes, indicating an association with CAR-T resistance. (b) Higher expression of a JAK/STAT/MHC-I resistance signature is associated with poor outcomes in CD19 CAR-T treated patients. (c) After CAR-T failure, relapsed B-ALL cells exhibit increased expression of immune checkpoint blockade resistance-associated interferon stimulated genes (ISG.RS) that is also associated with poor outcomes in CD19 CAR-T treated large B-cell lymphoma patients. (d) A final model for how high IFNγ/JAK/STAT signaling in tumor cells can promote resistance to CAR-T therapy via the upregulation of the inhibitory molecule Qa-1b, the murine homolog of HLA-E. Re-sensitization of tumor cells to CAR-T therapy can be accomplished by concurrent NKG2A blockade. Significance is determined using unpaired two-sided student’s t-tests. Data are mean ± s.e.m. *P<0.05; **P < 0.01; ***P < 0.001; ****P < 0.001.

## Discussion

Despite unprecedented success in treating B-cell malignancies, a majority of patients who receive CAR-T therapy ultimately relapse, highlighting the critical need to understand resistance mechanisms in order to improve clinical efficacy.^5, 8–13^ Taking a novel screening approach in an immunocompetent B-ALL model, we identified IFNγ/JAK/STAT signaling and components of the antigen processing and presentation machinery as critical *in vivo* mediators of CAR-T resistance in B-ALL. Here, loss of IFNγ/JAK/STAT pathway members or Qa-1^b^ rendered tumor cells more sensitive to CAR-T therapy *in vivo*. Paradoxically, loss of these pathways *in vitro* resulted in an opposite phenotype, or no phenotype at all, underscoring the importance of assaying CAR-T function in immunocompetent models of disease. This importance is augmented by the fact that our data, as well as that of other groups, suggests the endogenous immune system aids in, or perhaps is even crucial for CAR-T efficacy. For example, a recent study in an immunocompetent murine model of GBM found that CAR-T cells induce endogenous immune cell activation necessary for full antitumor efficacy.^55^ In fact, successful treatment with CAR-T cells has been shown to protect against subsequent challenges with antigen negative tumors, demonstrating the profound effect CAR-T cells can exert on the endogenous immune system.^56–58^ Notably, these results appear to be clinically relevant as studies in lymphoma and GBM patients have shown that CAR-T cells activate host immune cells and can even incite endogenous tumor specific T-cell responses.^59–61^ Thus, while the use of immunodeficient models is a predominant practice in the CAR-T field, our results suggest that CAR-T cell functionality should also be examined in the context of an intact immune system where crosstalk with the tumor microenvironment (TME), and the effects of this interaction can be appreciated.

Differences in the contexts in which CAR-T cells are used could help to explain apparent discrepancies between our results and those of other groups, as it relates to the role of IFNγ. For example, we show that global inhibition of IFNγ *in vivo* blocks CAR-T efficacy in the context of an immunocompetent B-ALL model, a finding that has been reproduced by other groups.^31, 33^ Interestingly, a recent study used immunocompromised and *in vitro* models to show that IFNγ is not required for CAR-T antitumor efficacy in the case of hematologic malignancies.^62^ However, it is well documented that the TME can negatively impact CAR-T cells, and that factors such IFN signaling can mediate both immune activation and suppression, depending on the cell type and whether signaling is acute or chronic. A recent salient example comes from CAR-T treated patients with large B cell lymphoma (LBCL). Here, Jain and colleagues find that high tumor IFN signaling and myeloid cells detected in pretreatment LBCL samples are associated with decreased CAR-T cell expansion and poor clinical outcomes.^52^ This IFN-stimulated gene signature (ISG.RS) was previously shown to be a marker of chronic IFN stimulation originating from tumor cells. In this context, malignant cells demonstrated increased surface expression of inhibitory T cell ligands, leading to T cell exhaustion and checkpoint therapy resistance.^63, 64^ Notably, when tumor IFN signaling was blocked and TME-based IFN signaling predominated, tumors could be rendered responsive to checkpoint blockade via reactivation of endogenous immune cells.^63^ Similarly, Jain and colleagues report that increased tumor expression of IFN controlled inhibitory ligands, such as PD-L1 and MHC-II, was associated with lack of durable treatment responses, raising the possibility that the same interplay between the TME, tumor cells, and CAR-T cells is at play in human malignancies. Thus, it is possible that in the context of the syngeneic tumors described in this study, production of IFNγ is necessary for treatment success, perhaps not because it helps kill tumor cells or acts directly on the CAR-T cells themselves, but instead because it recruits and activates endogenous immunity.

While our screen highlighted the importance of IFNγ/JAK/STAT signaling in promoting resistance to CAR-T therapy in B-ALL, additional data suggested that there may be no therapeutic window for global IFNγ blockade or JAK2 inhibition combined with CAR-T therapy. Importantly, our follow up validation screen allowed us to hone on in on Qa-1^b^ (mouse homolog of human HLA-E), a downstream target of IFNγ. B-ALL cells unable to signal through the IFNγ/JAK/STAT pathway were functionally deficient for Qa-1^b^ suggesting that the mechanism by which IFNγ/JAK/STAT signaling promotes CAR-T resistance in this context is, at least in part, dependent on IFNγ-induced cell surface expression of Qa-1^b^. Furthermore, we observed that CAR-T efficacy was enhanced by blocking NKG2A, the inhibitory receptor of Qa-1^b^, present on NK cells and subsets of T cells. Interestingly, NKG2A has been implicated as a novel immune checkpoint protein and blocking antibodies against it have been shown to have beneficial antitumor effects by acting on endogenous NK and T cell populations.^36, 43, 65, 66^ While these studies have focused on the application of this antibody alone, our data suggests that CAR-T therapy may also benefit from combination with this blocking antibody. At present, the mechanism behind this benefit is unclear and, interestingly, may not involve endogenous NK cells. Surprisingly, the addition of an NK1.1 blocking antibody, commonly used to deplete NK cells *in vivo*, improved CAR-T efficacy. Taken together, our data suggest that while the full functionality of CAR-T cells ostensibly depends on the activation of endogenous immune cells, the impact of the Qa-1^b^/NKG2A interaction is likely most significant between CAR-T and tumor cells in terms of imparting therapeutic resistance. Beyond this, IFNγ acts as a two-edged sword in that it induces expression of Qa-1^b^ (and potentially other inhibitory proteins) on tumor cells but also potentially aids in recruiting or activating endogenous immunity that is ultimately necessary for CAR-T antitumor efficacy. It remains to be determined the extent to which this finding applies in other tumor contexts and whether CAR-T treatment can benefit from the addition of NKG2A blocking antibody in the context of other malignancies.

Apart from therapeutic implications, our results also have potential for use in patient stratification for CAR-T therapy. Using published patient data, we showed that elevated expression of a JAK/STAT/MHC-I signature in pre-treated patient samples was associated with poor outcomes. Thus, certain tumors may possess transcriptional profiles rendering them more resistant to CAR-T therapy from the beginning. Examination of additional data sets from patients receiving CAR-T therapy will be required to test this hypothesis. Ultimately, our data highlight new strategies for both the design of improved CAR-T treatment regimens and for the development of pre-treatment tumor expression profiles that may help predict which patients are most likely to respond.

Lastly, a number of groups have employed screening-based functional genomics approaches to interrogate tumor intrinsic mechanisms of CAR-T killing however, these have all been conducted *in vitro*. Our screen is the first to ever be performed *in vivo*. The generation of our novel, small-pooled sgRNA library, along with our follow up validation screening approach, were critical for completing this study. To generate truly robust *in vivo* screening data, it is necessary to determine tumor engraftment rates and subsequently integrate these cancer model limitations when determining screening parameters. The value in our custom library is ultimately its modularity, allowing for use in a broad range of *in vivo* tumor settings. Most importantly, given the complexity of the tumor microenvironment and the interplay between endogenous immunity and immunotherapies like CAR-T, we believe conducting *in vivo* screens can provide valuable insights to improve therapies that would not have been uncovered otherwise.

## Methods

### Pooled sgRNA screening

A custom genome-wide library divided into 48 sub pools was generated in collaboration with the Broad Institute’s Genetic Perturbation Platform (GPP). In total, 97,336 unique guides targeting the protein coding regions of 21,958 unique murine genes with 4 sgRNAs each (plus control non-targeting and intragenic cutting guides) were included. All protein coding murine genes were subdivided into 48 pools by their initial KEGG term (obtained using KEGG REST API in BioPython, biopython.org), in a non-redundant manner. All four guides targeting the protein coding region of any given gene were kept together in the same pool. Using this approach, only 36% of protein-coding genes could be classified into a KEGG pathway. Thus, the first 14 sub pools and part of sub pool 15 were filled by KEGG genes. All other genes were randomly distributed among the remaining sub pools. Mouse essential genes (defined as orthologous mouse genes for the human essential gene set from Hart et al., 2015, 1530 genes; obtained from Ensembl Biomart) were divided evenly across all pools.^67^ Guides against human *EGFRvIII*, human *CD22*, human *CD19*, and murine *Cd19* were included in the first pool. Guides against olfactory genes (1,133 total) were also distributed evenly amongst all sub pools. All 48 sub pools were cloned into a lentiviral pRDA-Crimson_170 vector (Figure 1o). To preserve library complexity, a minimum of 1000-fold coverage of the sgRNA library was maintained at each *in vitro* step before the screen, and at a minimum of 150-fold coverage (range: 153 to 203-fold coverage *in vivo*, all *in vitro* screens were performed above 500x) was maintained in all screens completed. Pool A consisted of the first 8 sub pools and had a total of 15,308 sgRNAs targeting the protein coding regions of 3,648 unique mouse genes. Pool B consisted of sub pools 9-16 and had a total of 15,147 sgRNAs targeting the protein coding regions of 3,648 unique mouse genes. Pool C consisted of sub pools 17-24 and had a total of 15,258 sgRNAs targeting the protein coding regions of 3,648 unique mouse genes. Pool D consisted of sub pools 25-32 and had a total of 17,335 sgRNAs targeting the protein coding regions of 3,648 unique mouse genes. Pool E consisted of sub pools 33-40 and had a total of 14,713 sgRNAs targeting the protein coding regions of 3,648 unique mouse genes. Pool F consisted of sub pools 41-48 and had a total of 19,575 sgRNAs targeting the protein coding regions of 3,718 unique mouse genes. Cloned and sequenced plasmid pools, and viral supernatant were generated by the Broad Institute’s GPP.

For screens, Cas9^+^ cells were thawed, recovered, and expanded for 5 days to ensure robust growth, and then tested for cutting efficiency to ensure high rates of editing efficiency.^27, 68^ After cutting assays were completed, cells were expanded over three additional days and infected with sub pools. For each of the 48 sub pools, 6×10^7^ cells were spin-infected with predetermined amounts of viral supernatant (determined using titration experiments, data not shown), such that 15-30% of all cells were infected (expressed E2-Crimson, and survived puromycin selection; MOI<<1). The resulting cells were resuspended at a concentration of 10^6^ cells/mL in virus-containing medium supplemented with 10 μg/mL polybrene (Sigma), divided into 6-well plates, and centrifuged at 1000xg and 37°C for 1.5 hrs. Cells were then pooled into flasks and cultured overnight. Thirty-six hours later, cell density was adjusted to 10^6^ cells/mL (and was never allowed to go over 3×10^6^ cells/mL) and puromycin selection (2 μg/mL, Gibco, A1113803) was started. Cells were selected over two days and then spun out of puromycin-containing medium. Cells were then allowed to recover for one day, after which the appropriate number of infected, selected, and recovered cells were flow cytometry sorted and combined into their respective pools. The next day, cells from large, combined pools were prepared for tail vein injection into mice or kept in culture for the *in vitro* screens. Two days later, CAR-T cells were adoptively transplanted into mice via tail vein injection at the indicated doses. For all *in vitro* screens, 1.4×10^7^ library cells were seeded and treated two days later (on the same schedule as mice) with control CAR-T cells (anti-hEGFRvIII), anti-mCD19 CAR-T cells, or with no CAR-T cells, at E:T ratios indicated in the text. *In vitro* CAR-T screens were set up in triplicate while the no CAR-T condition was kept in a single plate. Input samples were collected just after puromycin selection had completed (Input PS) and on the day cells were injected into mice/set up for *in vitro* screens (Input or Input DOI). Upon becoming moribund, mice were sacrificed and E2-Crimson^+^ cells were sorted from the bone marrow and spleen (average number sorted cells per compartment = 1.38×10^7^). For *in vitro* screens where the B-ALL cells viability was above 95% and represented more than 99% of all cells present in the sample, cells were counted and 2×10^7^ cells were collected for gDNA isolation. The only condition that did not meet this cutoff was the anti-mCD19 at E:T of 1:2 for all 6 pools. Here, CAR-T cells were completely gone from culture, but B-ALL viability of cells remained low at 30-60% across replicates, necessitating sorting to isolate cells (1.4-2.0×10^7^ sorted per replicate, per condition). For all flow cytometry experiments in animal samples (screens and validation), data for a minimum of 10,000 live cells (via DAPI exclusion) were collected and analyzed.

Finally, gDNA from all cells were isolated using the Machery Nagel L Midi NucleoSpin Blood Kit (Clontech, 740954.20). Modifications to the manufacturer’s instructions were added as follows: in step 1, cells were lysed in the kit’s proteinase K containing lysis buffer for longer (overnight at 70°C). The next morning, lysates were allowed to cool to room temperature, 4.1 μL of RNase A (20 mg/mL; Clontech, 740505) was added, and cells were incubated for 5 minutes at room temperature. The procedure then continued as indicated by the manufacturer. PCR inhibitors were removed from the resulting gDNA (Zymo Research, D6030) and the concentration of the resulting gDNA was measured using the Qubit dsDNA HS assay kit (ThermoFisher, Q32854), and if necessary, diluted to 200 ng/μL with elution buffer. gDNA was then submitted for Illumina sequencing.

### Screen hit discovery

Sample quality control was performed by counting the number of sgRNA sequences that show at least 30 reads in each sample. Samples with more than 30% of sgRNA sequences that were filtered out after the above procedure were excluded from further analyses. Guide RNA-level read counts were scaled by total read count for each sample and logarithm-transformed. Gene-level enrichment and depletion scores were computed by averaging log-normalized read counts across all sgRNA sequences against each gene. Six different pooled libraries were aggregated targeting a total of 21,958 unique murine genes with a total of 88,793 sgRNAs (4 per gene plus non-targeting and intergenic-cutting controls). Input samples were used as a baseline for computing scores. Specifically, every gene was first assigned an ‘essentiality score’ computed as the difference (i.e., log-fold change) between the average gene-level counts for the gene across replicates and that in the input sample. The CAR-T therapy enrichment/depletion score for a given gene is computed as the difference between the essentiality scores of that gene in anti-mCD19 CAR-T cell treated samples and in anti-hEGFRvIII CAR-T cell treated samples. For sample groups that have at least 2 samples retained after quality control, Student’s t-tests were performed to obtain p-values for assessing significance of difference between the scores in the treatment group and the control group.

### Bulk transcriptome profiling and analysis

Total RNA was extracted from 1×10^6^ cells per sample using the Macherey-Nagel Nucleospin RNA Plus kit, and RNA sample quantity and quality was confirmed using an Agilent Fragment Analyzer. RNAseq libraries were created from 250ng of total RNA using the NEBNext UltraII Directional RNA Library Prep kit (New England Biolabs) using half volume reagents and 14 cycles of PCR. Illumina library quality was confirmed using the Fragment Analyzer and qPCR and sequenced on an Illumina NextSeq500 using v2.1 chemistry with 40nt paired end reads (RTA version 2.11.3). Paired-end RNA-seq data was used to quantify transcripts from the mm10 mouse assembly with the Ensembl version 101 annotation using Salmon version 1.3.0.^69^ Gene level summaries were prepared using tximport version 1.18.0^70^ running under R version 4.0.3.^71^

### Single cell transcriptome profiling and data processing

Mice were transplanted with 3×10^6^ B-ALL cells and challenged with 10^7^ CAR-T cells targeting either mCD19 or a control epitope (human EGFRvIII). Upon becoming moribund, mice were sacrificed, and B-ALL cells were sorted from the bone marrow and spleen. For each sample, approximately 10,000 live B-ALL cells (via DAPI exclusion) were sorted for transcriptional profiling. Single-cell expression libraries were prepared using the 10x Genomics Chromium v3 reagents. Sequencing data was aligned to the mm10 reference genome and converted to fastq files using bcl2fastq (v2.20.0.422). Cell count matrices were generated using cellranger (v.5.0). Matrices were analyzed by Seurat (v4.0.4) for R (v4.0.2). Digital gene expression matrices were filtered to exclude low quality cells (< 1000 UMI, < 400 genes or > 8000 genes, > 50% mitochondrial reads). Low-quality cells were further filtered from the dataset using the variance sink method as previously described.^72^ Briefly, data was normalized and scaled, known cell cycle genes were regressed out.^73^ Principal component analysis was performed on regressed and scaled data. Standard deviation of principal components was quantified using an elbow plot, input dimensions for SNN clustering (EGFRv3 bone marrow = 35, EGFRv3 spleen = 42, CD19 bone marrow = 30, CD19 spleen = 30) at which standard deviation = 2. SNN clustering was performed to generate UMAP plots (k.param = 40, res = 0.5). Clusters containing low quality cells (50% of cells with > 10% mitochondrial reads) were removed from the dataset. After filtering, samples from bone marrow and spleen treated with EGFRv3-CAR-T or CD19-CAR-T were merged into a single dataset. Cell cycle phase was assigned using the cell cycle scoring function based on expression of known cell cycle genes.^73^ Merged data set was normalized and scaled, cell cycle genes were regressed out. Principal component analysis was performed, and standard deviation of principal components was quantified by elbow plot. Nearest neighbors were found (dim = 50, k.param = 40) then clustered using SNN clustering (res = 0.5). Enriched genes for each cluster were identified with the cluster marker function. Cluster occupancy was quantified for each treatment condition and phase of the cell cycle to further define therapeutic response.

### Mouse maintenance and studies

All mouse experiments were conducted under IUCAC-approved animal protocols at the Massachusetts Institute of Technology. The mouse strains used in this study included C57BL/6 (Jackson) and NOD/SCID/IL2Rg^−/−^ (NSG; Jackson Laboratory). Immunocompetent recipient mice were sub-lethally irradiated (1 x 5 Gy) immediately prior to transplantation with B-ALL cells, receiving ACT of CAR-T cells 2 days later, as noted in the text. For *in vivo* screens, mice were injected with 3×10^6^ Cas9^+^ library B-ALL cells and the indicated number of CAR-T cells. Both B-ALL and CAR-T cells were prepared for transplantation by resuspending in 200 μL Hank’s balanced salt solution (Lonza) and loaded in 27.5-gauge syringes (Becton Dickinson). All cell solutions were administered via tail vein injections. For validation experiments conducted with the murine glioblastoma line GL261, 0.5×10^6^ GL261 cells were delivered via intracranial injection into female C57BL/6 mice. Mice were then treated 4 days later with an intracranial injection of 0.2×10^6^ CAR-T cells. Surgical procedure closely followed that of previous studies using this model conducted in our lab.^74^ For *in vivo* blocking of IFNγ, mice were injected i.p. every third day with 200μg of InVivoMAb anti-mouse IFNγ antibody (clone XMG1.2, Bio X Cell) or 200μg of InVivoMAb anti-horseradish peroxidase control antibody (clone HRPN, Bio X Cell), starting the day after ACT. For *in vivo* inhibition of JAK1/2, Ruxolitinib (Selleckchem, INCB018424) was resuspended according to manufacturer guidelines and mice were dosed every twelve hours via oral gavage with 90 mg/kg Ruxolitinib or vehicle control, starting the day after disease transplantation. For *in vivo* blocking of NK1.1, mice were injected i.p. on the same day as ACT, and then once again 7 days later, with 200μg of InVivoMAb anti-mouse NK1.1 antibody (clone PK136, Bio X Cell) or 200μg of InVivoMAb mouse IgG2a isotype control antibody (clone C1.18.4, Bio X Cell). For *in vivo* blocking of NKG2A, mice were tail vein injected with 200μg of InVivoMAb anti-mouse NKG2A antibody (clone 20D5, Bio X Cell) or 200μg of InVivoMAb rat IgG2a isotype control antibody against trinitrophenol (clone 2A3, Bio X Cell) on the same day as ACT. Mice were re-dosed with their respective antibody three and six days later and monitored for survival.

### Bioluminescence studies

XenoLight D-Luciferin Potassium Salt D (PerkinElmer, 122799) was used for standard bioluminescent imaging (resuspended at 30 mg/mL in saline, sterile filtered, and stored at −80°C). Mice were weighed and luciferin was loaded in 27.5-gauge syringes (Becton Dickinson) and administered via intraperitoneal injection at a dose of 165 mg/kg. Mice were then anesthetized with 2.5% isoflurane (Piramal Critical Care, NDC# 66794-013-25), delivered at 1 L per minute in O_2_. Ten minutes from the time of luciferin injection, animals were imaged on a Xenogen IVIS system at consistent exposures between groups with small binning. Data was analyzed using Living Image version 4.4 software (Caliper Life Sciences). Images were normalized to the same color scale for figure generation.

### Cell culture

All cell lines were mycoplasma negative.

<u>Murine B-ALL cells:</u> Cells were cultured in RPMI with L-glutamine (Corning, 10-040-CM), supplemented with 10% fetal bovine serum (FBS) and 2-mercaptoethanol to a final concentration of 0.05mM (Gibco, 21985023).

<u>Murine B-cell lymphoma cells (Eμ-Myc):</u> Cells were cultured in medium composed of a 50:50 mix of IMDM with L-glutamine and 25mM HEPES (Corning, 10-016-CM) and DMEM with L-glutamine and sodium pyruvate (Corning, 10-013-CM), supplemented with 10% FBS and 2-mercaptoethanol to a final concentration of 0.05mM (Gibco, 21985023).

<u>Murine Glioblastoma cells (Gl261):</u> Cells were cultured in DMEM with L-glutamine and sodium pyruvate (Corning, 10-013-CM) supplemented with 10% FBS.

<u>Murine T cells:</u> T-cells harvested from the spleens of mice were cultured in plates coated with activating antibodies (as described in CAR-T cell production methods) in T-cell medium (TCM): RPMI with L-glutamine (Corning, 10-040-CM), supplemented with 10% FBS, recombinant human IL-2 (rhIL-2, final concentration of 20 ng/mL; Peprotech, Cat# 200-02-1mg), and 2-mercaptoethanol to a final concentration of 0.05mM (Gibco, 21985023).

<u>Human cell lines</u>: 293T and Raji cells were all mycoplasma negative. 293T cells were cultured in DMEM with L-glutamine and sodium pyruvate (Corning, 10-013-CM) supplemented with 10% FBS. Raji cells were cultured in RPMI with L-glutamine (Corning, 10-040-CM) supplemented with 10% FBS.

### Viral supernatant production

Viral supernatant was produced using standard methods. Briefly, 293T cells were transfected with retroviral or lentiviral transfer plasmid and packaging vector (retrovirus: pCL-Eco, Addgene, 12371; lentivirus: psPAX2, Addgene, 12260 with VSVg envelop plasmid pMD2.G, Addgene, 12259) using Mirus TransIT-LT1 (Mirus, MIR2305) as indicated by the manufacturer. The next day, 293T cells were switched into medium composed of 60% RPMI complete and 40% DMEM complete. Viral supernatant was collected 24 and 48 hours after transfection, passed through a 0.45um filter to remove residual 293T cells, and kept at 4°C for a maximum of 4 days.

### *In vitro* killing assay

*In vitro* CAR-T killing assays were performed using standard methods.^75, 76^ Briefly, target cells are counted and co-cultured with or without CAR-T cells at indicated E:T ratios (accounting for CAR-T infection rate) in RPMI complete supplemented with 2-mercaptoethanol and 10% FBS but no rhIL-2. Sixteen to twenty-four hours later, the total cell number per well was counted and the cell suspension was analyzed by flow cytometry to assess for live/dead (via DAPI stain), %mCD19^+^ cells, and %CD8^+^ cells. The densities of each cell type (CAR-T, target cell, non-transduced T cell) were also determined via flow cytometry. The resulting target cell densities in CAR-T containing wells are then normalized to the resulting target cell density in control wells seeded with the same number of target cells but without CAR-T cells. For all flow cytometry experiments, data for a minimum of 10,000 live cells (via DAPI exclusion) were collected and analyzed.

### Interferon gamma ELISA release assay

Standard methods were used for enzyme-linked immunosorbent assay (ELISA). Briefly, supernatant from *in vitro* CAR-T killing assays was collected and spun down to remove any contaminating cells. IFNγ released into the supernatant by CAR-T cells was then measured using the DuoSet ELISA kit for mouse IFNγ (R&D systems, DY485) and Nunc MaxiSorp flat bottom plates (Thermo Fisher Scientific, 44-2404-21) on a Tecan infinite 200 Pro machine, as indicated by the manufacturer. To ensure that the assay was completed within the linear range of the kit, supernatant is initially diluted 1:10 in reagent diluent. At least six serial 4-fold dilutions were then performed. At least one standard curve for this assay was generated per plate and at least two standard curves for the entire experiment were constructed using standard solutions supplied by the manufacturer. The substrate solution was 1-Step^TM^ Ultra TMB-ELISA (Thermo Fisher Scientific, 34028) and the stop solution was 2N sulfuric acid (VWR, BDH7500-1). Bovine serum albumin (BSA; Sigma, A8022-500G) was prepared as a sterile filtered 5% stock in PBS (Corning, 21-031-CV).

### CAR-T cell production

Before collecting T-cells, 6-well plates were coated overnight with activating antibodies against mCD3e (Bio X-Cell, BE0001-1) and mCD28 (Bio X-Cell, BE0015-1) at 5 μg/mL each in PBS (Corning, 21-031-CV) at 4°C. The next day, 8–12-week-old male C57BL/6 mice (Jackson) were sacrificed, and their spleens were collected. CD8^+^ T-cells were isolated using Miltenyi Biotec CD8a (Ly-2) MicroBeads for mouse (positive selection kit; Miltenyi, 130-117-044) and LS columns (Miltenyi, Cat# 130-042-401) as indicated by the manufacturer. Coated plates were rinsed once with PBS and T-cells were resuspended at 0.5×10^6^ to 10^6^ cells/mL in T-cell medium (TCM, recipe in cell culture methods). After 24 hours, activated T-cells were collected and placed into fresh TCM after counting. Cell concentrations were then adjusted to 10^6^ cells/mL in a 50:50 mix of TCM:retroviral (RV) supernatant supplemented with protamine sulfate to a final concentration of 10 μg/mL (MS Biomedicals, ICN19472910), 2-mercaptoethanol to a final concentration of 0.05 mM, and rhIL-2 to a final concentration of 20 ng/mL. Once resuspended, cells were spin-infected at 1000xg for 1.5 hrs at 37°C on new antibody coated plates. The next day, T-cells were collected from plates, resuspended in fresh TCM at a cell density of 0.5×10^6^ to 10^6^ cells/mL, and re-plated on new antibody-coated plates. Twenty-four hours later, T-cells were collected from antibody coated plates and prepared for tail vein injection into animals or for *in vitro* kill assays/screens, as described above. T-cells were always cultured and infected on PBS rinsed, antibody-coated 6-well plates, as described above, except during *in vitro* killing assays where no activating antibodies were ever used.

### Western Blotting

Cells were lysed with RIPA buffer (Boston BioProducts, BP-115) supplemented with 1X protease inhibitor mix (cOmplete EDTA-free, 11873580001, Roche). Protein concentration of cell lysates was determined using Pierce BCA Protein Assay (ThermoFisher Scientific, 23225). Total protein (40-60μg) was separated on 4-12% Bis-Tris gradient SDS-PAGE gels (Life Technologies) and then transferred to PVDF membranes (IPVH00010, EMD Millipore) for blotting.

### Plasmids, cloning, and sgRNAs

#### Packaging and envelope plasmids used for viral production

Retrovirus: pCL-Eco (Addgene, 12371)

Lentivirus: psPAX2 (Addgene, 12260) with VSVg envelop plasmid pMD2.G (Addgene, 12259) or pCMV-EcoEnv (Addgene, 15802)

#### Chimeric Antigen Receptor (CAR) plasmids

The murine CD19 targeting second generation CAR 1D3-28Z.1-3 containing inactivating mutations in the 1^st^ and 3^rd^ ITAM regions of the CD3-ζ chain was synthesized by Twist Bioscience and cloned into the GFP^+^ MP71 retroviral vector.^28, 77^ The clinically used scFv sequence (heavy chain linked to light chain variable regions) against human CD19, FMC63, was provided by the Maus lab. A CD28-containing 2^nd^ generation murine CAR targeting hCD19 protein was then constructed by switching out the scFv for 1D3-28Z.1-3 in the anti-mCD19 CAR and replacing it with the FMC63 scFv (Twist Bioscience). The same technique was used for the 3C10 scFv targeting human EGFRvIII, which was reported by the Rosenberg lab.^78^ All CAR constructs are identical, containing a CD8a leader sequence, followed by the respective scFv, followed by an IgG4 hinge sequence, a portion of the murine CD28 molecule from amino acids IEFMY to the 3′ terminus, and finally, the cytoplasmic region of the murine CD3-ζ chain from amino acids RAKFS to the 3′ terminus with both tyrosines in ITAMs 1 and 3 mutated to phenylalanines as described (all in frame).^28^ All CAR constructs were extensively tested to ensure that they only targeted their peptide of interest. Human EGFRvIII expression was induced using pMSCV-XZ066-EGFRvIII (Addgene, plasmid 20737) and murine hEGFRvIII^+^ B-ALL cells were generated. Retroviral supernatant to induce hCD19 expression was provided by the Maus lab. All CARs were extensively tested *in vitro* (killing assays methods) and *in vivo* (as described in text) to ensure no off-target effects were present.

#### CRISPR plasmids

To generate Cas9^+^ murine B-ALL cell lines, lentiCas9-Blast (Addgene, 52962) was used, and cells were selected with Blasticidin (Gibco, A1113903) at 20 μg/mL for 7 days and then single cell cloned and assayed for Cas9 expression via WB. Guide RNAs for murine *Cd19* were designed using the Broad Institute’s sgRNA Designer tool (https://portals.broadinstitute.org/gppx/crispick/public) and cloned into lentiGuide-Puro (Addgene, 52963) for the functional cut assay (tracking loss of mCD19 on the cell surface) or pRDA-Crimson_170 to generate KOs of various test genes (vector testing, data not shown).^27^ Guide RNAs used are below (Forward/Reverse):

#### Murine *Cd19* sgRNAs

sgRNA#42, targets exon 6 (5’-CACCGAATGACTGACCCCGCCAGG)/(5’-AAACCCTGGCGGGGTCAGTCATTC)

sgRNA#43, targets exon 2 (5’-CACCGCAATGTCTCAGACCATATGG)/(5’-AAACCCATATGGTCTGAGACATTGC)

#### Other plasmids

MSCV-mCherry (Addgene, 52114) was used to generate 20.12DP cells from mCherry^−^ GFP^+^ 20.12 cells.

### Generating Qa-1^b^ knockout B-ALL cells

B-ALL cell lines deficient for Qa-1^b^ were generated using CRISPR-Cas9 technology. Single guide RNAs directed against the *H2-T23* locus were designed using the Broad Institute’s sgRNA Designer tool and the sgRNA sequence (5’-GTACTACAATCAGAGTAACGA-3’) was cloned into pRDA-Crimson_170, as described above. Cas9^+^ B-ALL cells (clone RH62) were transduced with sgRNAs and selected with puromycin over 48 hours. Loss of Qa-1^b^ was confirmed by incubating transduced cells with 30 IU/mL IFNγ for 24 hours and subsequently analyzing Qa-1^b^ surface expression by flow cytometry, as previously described.^43^ Qa-1b^−^ cells were then FACS sorted twice until a pure population of Qa-1^b^ knockout cells was established.

### Antibodies

Western blotting: anti-β-ACTIN (Cell signaling, 4967S), anti-CD19 (Abcam, ab25232), anti-Cas9 (ActiveMotif, 61577), anti-JAK2 (Cell Signaling Technology, 3230), anti-IFNγR1/CD119 (R&D Systems, MAB10261), anti-STAT1 (Cell Signaling Technology, 9172), anti-rabbit IgG HRP-linked antibody (Cell Signaling Technology, 7074), anti-mouse IgG HRP-linked antibody (Cell Signaling Technology, 7076), anti-rat IgG HRP-linked antibody (Cell Signaling Technology, 7077), Rabbit anti-Armenian Hamster IgG H&L (HRP) (Abcam, ab5745).

Flow cytometry: anti-mouse CD19-BV785 (BioLegend, 115543), anti-mouse CD8-PE/Cy7 (BioLegend, 100722), anti-human CD19-APC/Cy7 (BioLegend, Cat# 302218), anti-mouse Qa-1^b^-BV786 (BD Biosciences, 744390).

## Statistical analysis

All Statistical analyses were performed using GraphPad Prism 9 (GraphPad Software Inc). The specific statistical tests performed are specified in figure legends. Differences are considered significant for P-values ≤ 0.05, or as indicated when adjustments for multiple hypothesis testing was required.

## Data Availability

RNA sequencing data generated in this manuscript is publicly available under GEO accession ID GSE196143. Sequencing data from all screens are available from the corresponding author on reasonable request. All other data are included within the manuscript figures, as supplementary information, or in supplementary table 1.

## Supporting information

Supplemental full wester blot figures

## Acknowledgements

The authors thank Hojun Li, and members of the Hemann and Vander Heiden labs for valuable discussions and intellectual input, the Koch Institute’s Robert A. Swanson (1969) Biotechnology Center for technical support, specifically the Flow Cytometry and Preclinical Imaging and Testing facilities, and S. Levine from the MIT BioMicro Center for informative discussions about guide library and scRNA sequencing. The authors also thank Beatrice Grauman-Boss for her support generating CAR virus, and Mudra Hegde for assistance with guide library design. This work was generously supported by the MIT Center for Precision Cancer Medicine, the Ludwig Center at MIT, a Margaret A. Cunningham Immune Mechanisms in Cancer Research Fellowship, a David H. Koch Graduate Fellowship, and the Paul and Daisy Soros Fellowship for New Americans. This work was also supported in part by NCI R01-CA233477, R01-CA226898 and NIH/NIAID R21AI151827 to M.T. Hemann and the Koch Institute Support (core) Grant P30-CA14051 from the NCI. The project described was supported by award Number T32GM007753 and T32GM144273 from the National Institute of General Medical Sciences. The content is solely the responsibility of the authors and does not necessarily represent the official views of the National Institute of General Medical Sciences or the National Institutes of Health

## Author information

Please note that Aviv Regev’s initials are noted below as AV to differentiate them from those of co-author Azucena Ramos.

## Contributions

AR, YL, and MTH conceived the idea for the study. AR, YL, CK, and MTH designed the study. AR, YL, CK, RH, KA, JF, DG, and KG conducted experiments. RH provided support with characterization of clone 20.12 and clone RH62, and the entirety of the primary screens. RCL provided support generating clone 20.12. TK and RCL provided critical support in designing the CAR constructs. TK cloned all CAR constructs. YL and DG analyzed scRNAseq data. AR and KA completed the validation screen. AV, JGD, MM, MVD, and MEB gave vital advice and provided reagents. AR, YL, CK, and MTH wrote the manuscript. All authors reviewed and edited the manuscript.

Corresponding author: Michael T. Hemann –

---

**Supplementary Figure 1.**
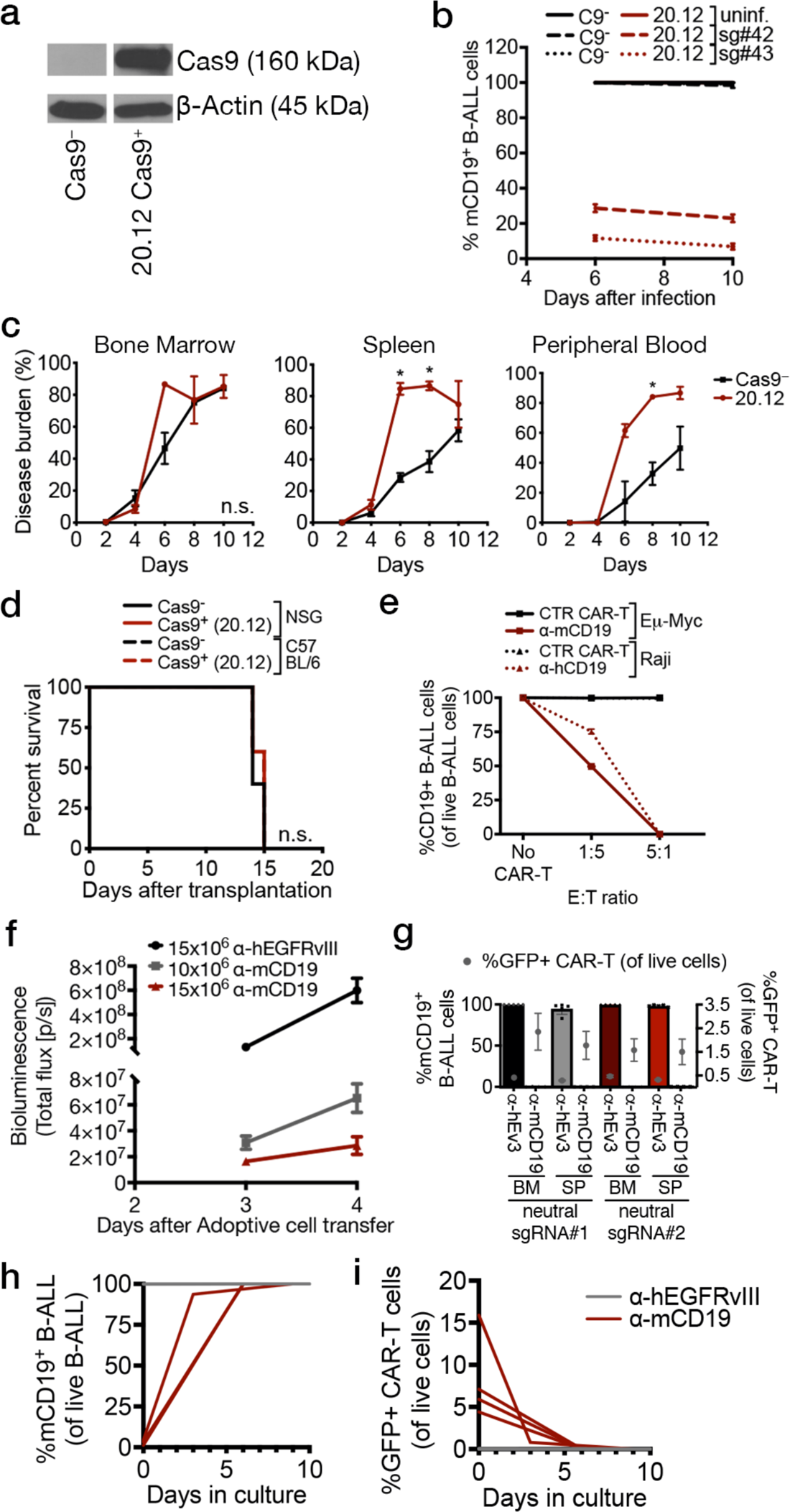
Characterization of Cas9+ disease and target epitope loss after CAR-T treatment. (a) Cas9 expression in 20.12 cells assayed via western blot. (b) *In vitro* cut assay using two validated, non-overlapping guides against murine CD19 (mCD19) was used to determine Cas9 cutting efficiency in 20.12 cells. Eight days after the introduction of sgRNAs, more than 75% of all 20.12 cells are mCD19^−^. (c) Tomato^+^ GFP^+^ Cas9^+^ 20.12 cells show similar or faster growth kinetics as non-Cas9 expressing parental cells in indicated hematopoietic organs of non-irradiated syngeneic B6 mice, as assayed by flow cytometry. (d) Non-irradiated immunocompetent (B6) mice and immunocompromised (NOD/SCID/IL-2Rγ, or NSG) mice succumbed to both Cas9^−^ and Cas9^+^ (20.12) disease at the same time, indicating that Cas9 is not additionally immunogenic *in vivo.* (e) Murine CAR-T cells targeting either mCD19 or human CD19 (targeted by the hybridoma FMC63 single chain variable fragment) *in vitro* induce dose dependent target epitope loss in murine Eµ-Myc and human Raji cells, respectively. (f) Bioluminescence imaging of irradiated B6 mice treated with indicated CAR-T cell type and dose at indicated time after ACT. On the day of ACT (two days after injection of B-ALL disease), disease burden is undetectable by bioluminescence imaging. (g) Organs from moribund control mice that relapsed after treatment with 3.5 ×10^6^ CAR-T cells targeting mCD19 demonstrate ongoing CAR-T engraftment and complete target epitope loss in relapsed B-ALL cells. The proportion of mCD19 positivity in live B-ALL cells harvested from relapsed animals treated with indicated CAR-T cell type is shown with bar graphs and the left sided y-axis. The proportion of live cells that are GFP^+^ CAR-T cells in relapsed animal organs is shown with grey dots and the right sided y-axis. (h-i) Whole spleens of relapsed mice after treatment with 5×10^6^ indicated CAR-T cell type were dissociated, cultured *in vitro* and serially assessed for mCD19^+^ B-ALL (h, showing proportion of live B-ALL cells that are also mCD19 positive) and GFP^+^ CAR-T cells (I, showing proportion of all live cells that are GFP^+^ CAR-T cells). As anti-mCD19 CAR-T cells are depleted out of culture (i, red lines), B-ALL cells re-express mCD19 (h, red lines). All experiments were repeated at least once. Significance for survival experiments was determined using log-rank tests. For all other experiments, significance is determined using unpaired two-sided student’s t-tests with Bonferroni correction for multiple comparisons. Data are mean ± s.e.m.; n = 4–8 mice per group. *P<0.05; **P < 0.01; ***P < 0.001; ****P < 0.001.

**Supplementary Figure 2.**
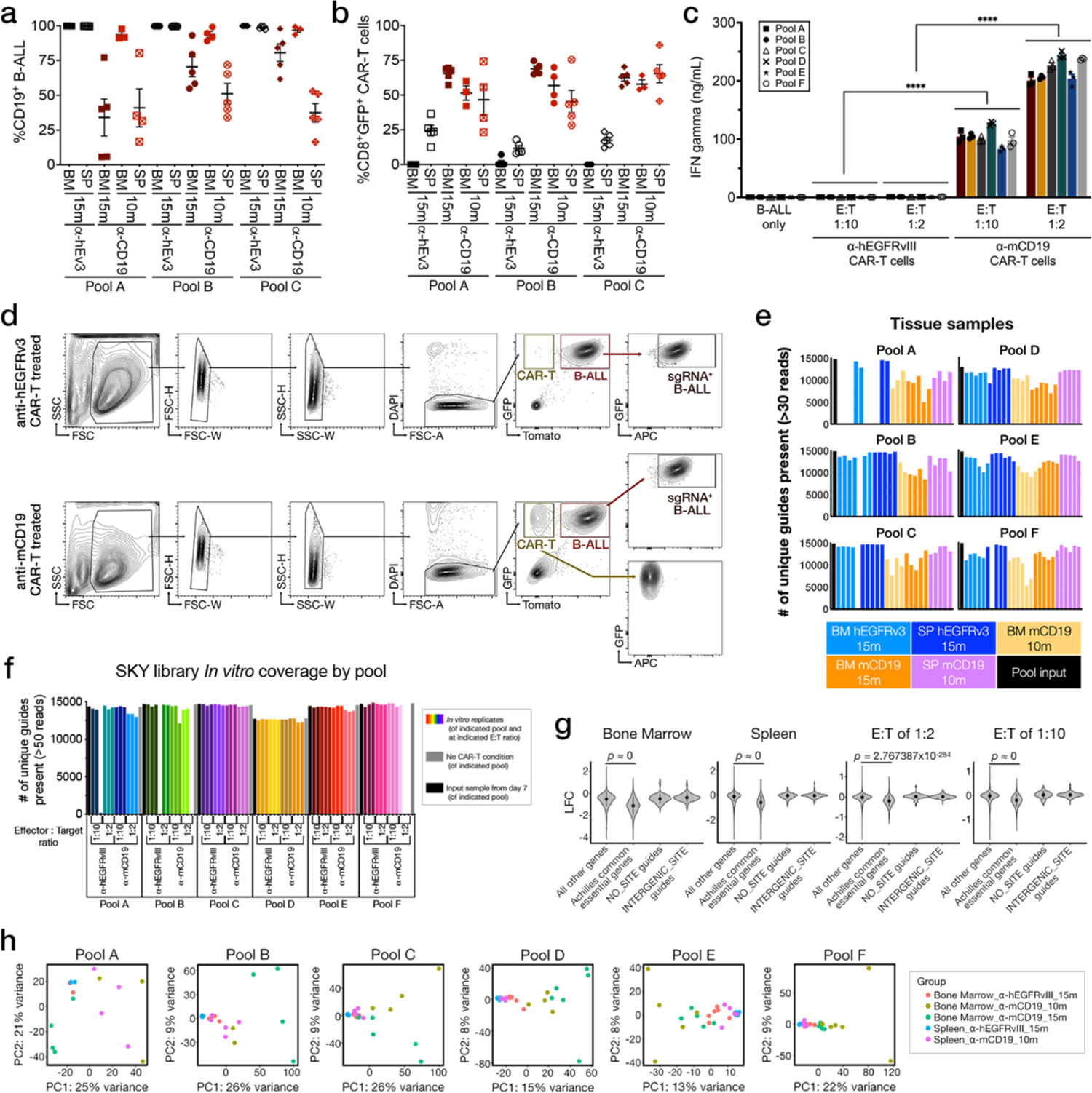
*In vitro* and *in vivo* genome-wide CRISPR/Cas9-mediated primary screens show comprehensive representation of sgRNA libraries and capture differences in sgRNA behavior in between control and anti-mCD19 CAR-T treated mice. (a) Flow cytometry assay investigating the proportion of live B-ALL cells that are mCD19^+^ in the spleen (SP) or bone marrow (BM) after treatment with the indicated CAR-T cell type (anti-mCD19 versus control anti-hEGFRvIII) and dose (10m and 15m indicate 10^7^ and 1.5×10^7^ CAR-T cells, respectively). (b) Flow cytometry assay investigating the proportion of GFP^+^ persistent CAR-T cells (amongst all live CD8+ T-cells) in the spleen (SP) or bone marrow (BM) after treatment with the indicated CAR-T cell type and dose. (c) IFNγ release quantified via ELISA in cell culture supernatant collected 24 hours after *in vitro* kill assays with murine B-ALL cells during primary screens. These experiments were completed in triplicate for each pool, each treatment type, and each E:T ratio (90 samples total). Anti-mCD19 CAR-T cells release significantly more IFNγ when exposed to their target antigen, as compared to control CAR-T cells. Differences between the two batches of CAR-T cells used in the primary screens (experiment 1: pools A-C; experiment 2: pools D-F). could not be detected using this assay. (d) Fluorescence activated cell sorting scheme used to isolate live, guide bearing (sgRNA^+^) B-ALL cells during both the primary and validation screens. (e) sgRNA library representation in each pooled library in the *in vivo* primary screen samples. Each bar shows the number of sgRNAs with more than 30 reads in the sequencing sample for a given pool. (f) Same as (e), but for sgRNA library representation in each pooled library in the *in vitro* primary screen samples. (g) Violin plots showing relative depletion/enrichment of control sgRNAs used in the primary screen. *P*-values are calculated from Wilcoxin rank-sum tests for comparison between pairs of group means. Note that in some cases exact *p*-values cannot be determined due to extremely near-zero values and ties in data and are reported as approximately zero. (h) *In vivo* primary screen samples projected onto the first 2 principal components of input-normalized sgRNA count matrix. The two experiments comprising the primary screen were completed one time each. For (a-c), data are mean ± s.e.m., n = 3–6 mice per group, and significance is determined using unpaired two-sided student’s t-tests with Bonferroni correction for multiple comparisons. *P<0.05; **P < 0.01; ***P < 0.001; ****P < 0.001.

**Supplementary Figure 3.**
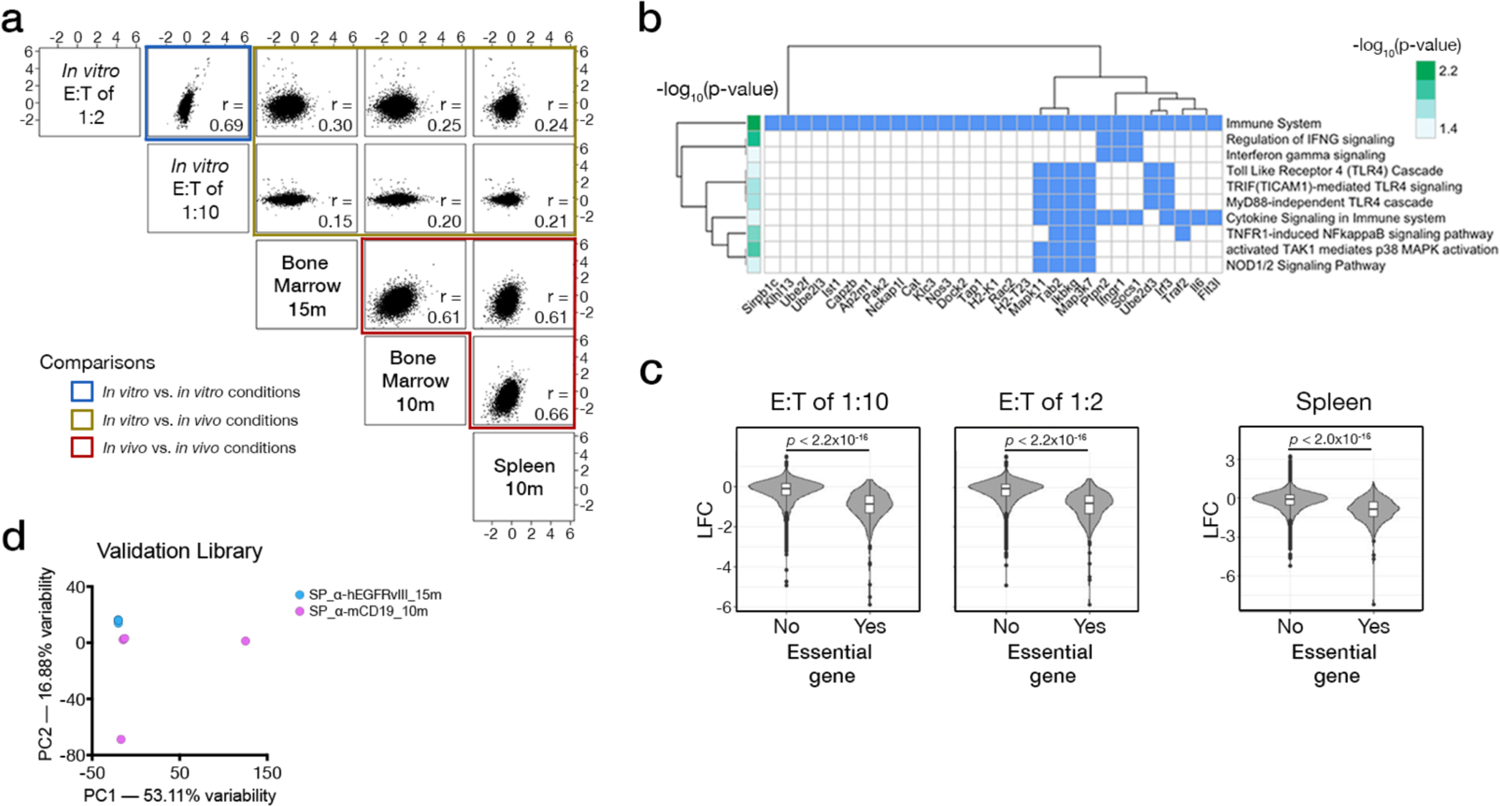
Both genome-wide and validation CRISPR/Cas9-mediated screens demonstrate robust quality control characteristics. (a) Pair-wise correlations between each arm of the primary screens. Top gene hits from both *in vitro* screen conditions are highly correlated with one another, demonstrating a Pearson correlation coefficient close to 1 (blue box). Top gene hits from all *in vivo* screen conditions are also highly correlated with one another, with Pearson correlation coefficients close to 1 (red box). However, in all cases, top hits from *in vitro* versus *in vivo* conditions are poorly correlated to one another and demonstrate weak Pearson correlation coefficients closer to 0 (gold box). (b) Enrichment of KEGG terms within genes showing top sgRNA depletion in the *in vivo* arms of the screen. Shown is a heatmap of participation of the top genes in each enriched pathway. Significance was calculated with hypergeometric tests corrected for multiple hypothesis testing to determine whether pathway terms are overrepresented among *in vivo* hits. (c) Violin plots showing relative depletion/enrichment of control sgRNAs used in the validation screen. *P*-values are calculated from Wilcoxin rank-sum tests for comparison between pairs of group means. (d) *In vivo* validation screen samples projected onto the first 2 principal components of input-normalized sgRNA count matrix. The two experiments comprising the primary screen were completed one time each. The validation library was also completed once. For the validation library, each *in vitro* screening condition was completed in triplicate and n = 4 mice per treatment group in the *in vivo* arm.

**Supplementary Figure 4.**
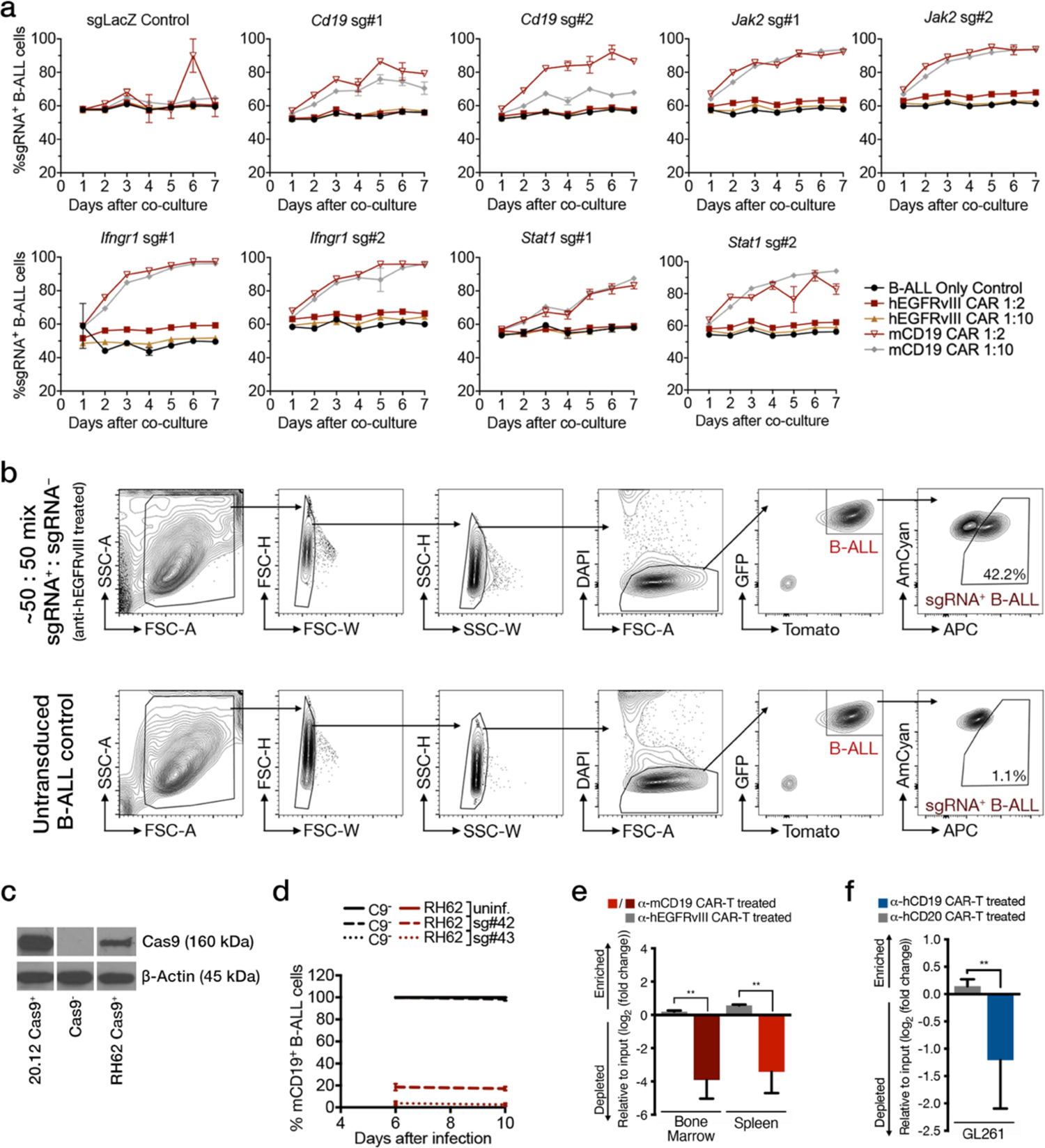
*In vitro* cytotoxicity assays in cells deficient for components of the IFNγ/JAK/STAT pathway do not capture *in vivo* sensitization phenotypes to CAR-T therapy. (a) *In vitro* competitive assays completed in cells transduced with guides against the indicated IFNγ/JAK/STAT pathway or control gene and treated with the indicated CAR-T cell type at two E:T ratios. Data shown is from flow cytometry analyses examining the proportion of live B-ALL cells that are APC^+^ and therefore, also guide-bearing. (b) Examples of the flow cytometry analysis scheme used for these experiments, along with untransduced B-ALL control cells used to guide gate placement. (c) Western blot showing that Tomato+ RH62 B-ALL cells used for all validation experiments express Cas9 protein. (d) *In vitro* cut assay using two validated, non-overlapping guides against murine CD19 (mCD19) was used to determine Cas9 cutting efficiency in RH62 cells. Six days after the introduction of sgRNAs, more than 75% of all RH62 cells are mCD19^−^, as assayed with flow cytometry. (e) *In vivo* competitive assays repeated in immunocompetent Cas9 transgenic mice again demonstrate specific log-fold depletion of Cas9^+^ *Ifngr1^—/—^* RH62 B-ALL cells after treatment with anti-mCD19 CAR-T cells in both the bone marrow (left) and the spleen (right). (f) *In vivo* competitive assays repeated in a murine syngeneic orthotopic transplant model of glioblastoma multiforme (GBM, GL261 cells) again demonstrate specific log-fold depletion of *Ifngr1^—/—^* GBM cells that express the extracellular domain of hCD19 after treatment with anti-hCD19 murine CAR-T cells. Anti-hCD20 murine CAR-T cells are used as controls. All *In vitro* experiments represented in this figure were repeated three times. *In vivo* experiments in (e) and (f) were completed once. Significance is determined using unpaired two-sided student’s t-tests with Bonferroni correction for multiple comparisons. Data are mean ± s.e.m.; n = 5 mice per group for *in vivo* experiments displayed. *P<0.05; **P < 0.01; ***P < 0.001; ****P < 0.001.

**Supplementary Figure 5.**
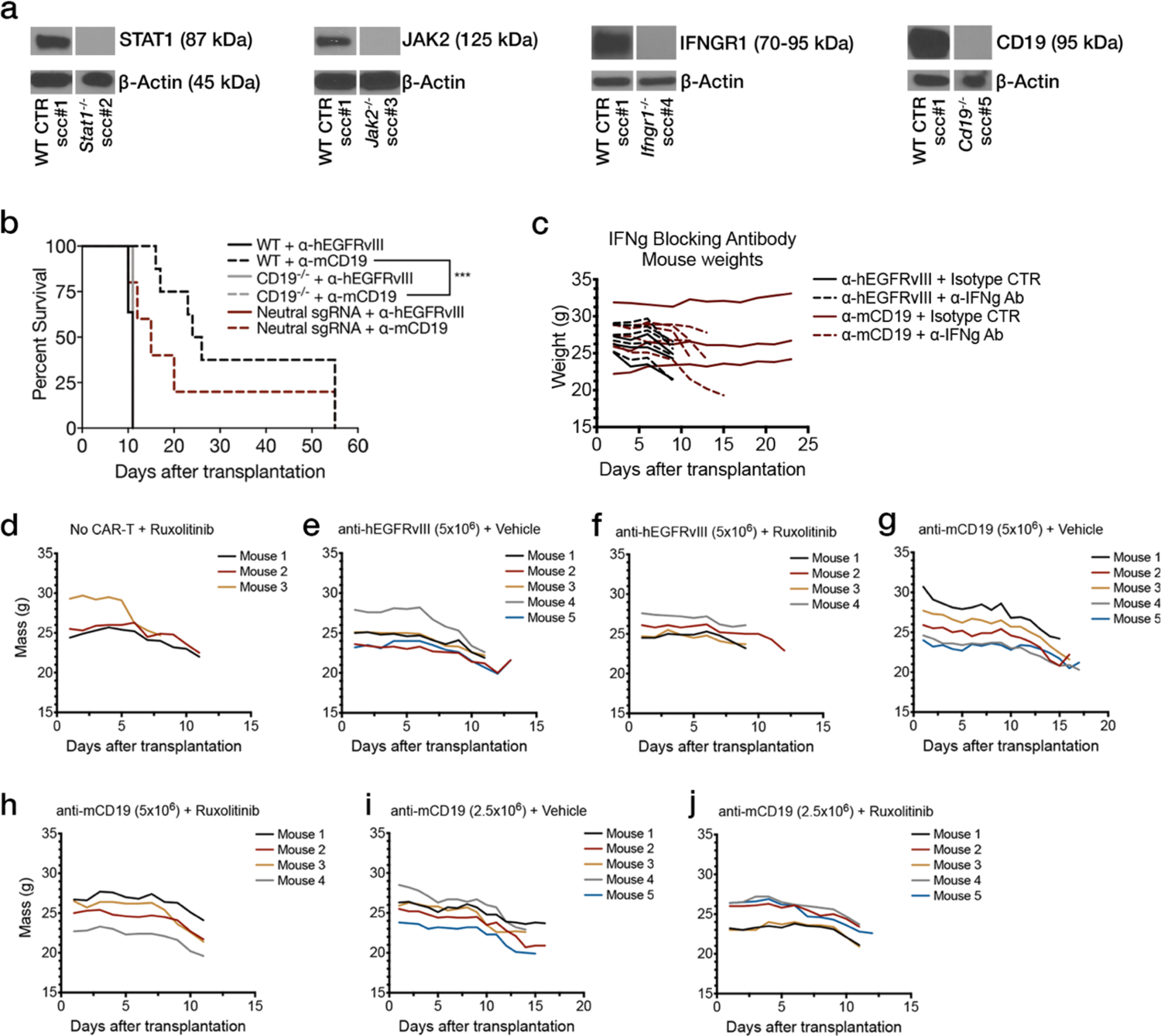
Generation of single cell B-ALL clones deficient in members of the IFNγ/JAK/STAT pathway and animal weight monitoring for pharmacological experiments targeting IFNγ/JAK/STAT signaling. (a) Western blot analysis of single cell clones (scc) deficient in indicated gene and used in survival validation experiments. (b) Mice transplanted with parental (WT) or neutral control cells do not significantly differ in their survival after treatment with anti-mCD19 CAR-T cells, while mice transplanted with *Cd19*^−/−^ B-ALL cells show no life extension. Here, n = 5 mice per group. (c) Mouse weight monitoring over time during co-treatment with CAR-T cells and anti-IFNγ blocking antibodies shows no additional toxicity during combination therapy. (d-j) Mouse weight monitoring over time during co-treatment with CAR-T cells and Ruxolitinib shows no additional toxicity during combination therapy. Data are exact mass in grams over time and the number of mice per group is indicated in each panel representing that group. Survival experiments were repeated three times. Pharmacologic experiments were completed once each. Significance for survival experiments was determined using log-rank tests. *P<0.05; **P < 0.01; ***P < 0.001; ****P < 0.001.

**Supplementary Figure 6.**
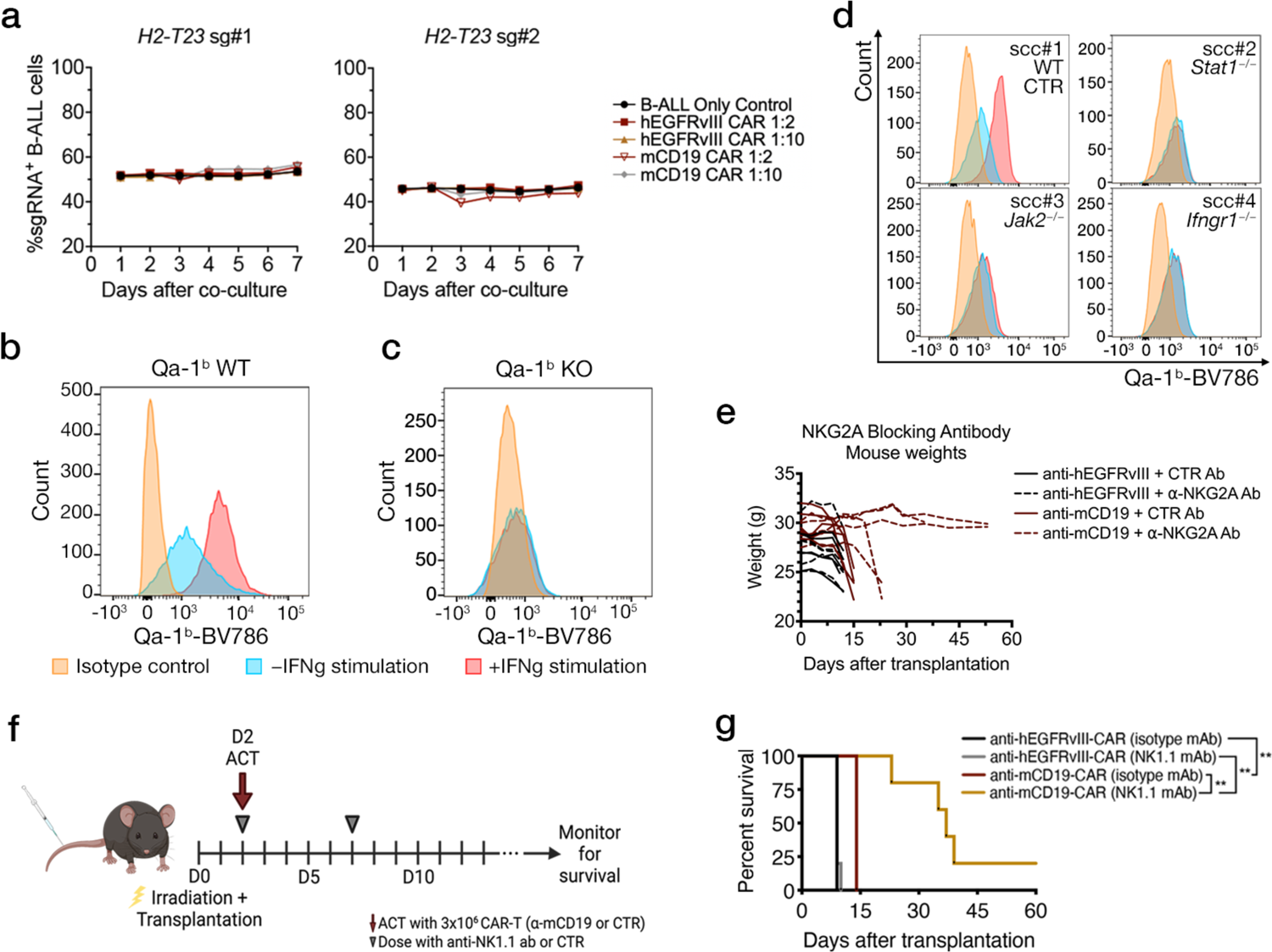
Loss of Qa-1^b^ does not sensitize B-ALL cells to CAR-T therapy *in vitro*. a) *In vitro* competitive assays completed in cells transduced with single guide RNAs (sg) against *H2-T23* and treated with the indicated CAR-T cell type at two E:T ratios. Controls for this experiment can be found in Supplementary figure 4a. Data are mean ± s.e.m. (b-c) FACS histograms showing Qa-1^b^ expression on cells transduced with a control sgRNA (Qa-1^b^ WT) or Qa-1^b^-deficient cells after stimulation with IFNγ. (d) Expression of Qa-1^b^-BV786 after *in vitro* stimulation with IFNγ of single cell clones (scc) deficient in indicated components of the IFNγ/JAK/STAT pathway, as assayed using flow cytometry. Wildtype (WT) control single cell clones were also generated and assayed at the same time. Representative data for a control WT clone is shown in (d). Color key is identical to that in (b, c). (e) Mouse weight monitoring over time during co-treatment with CAR-T cells and NKG2A blocking antibody shows no additional toxicity during combination therapy. Data are exact mass in grams over time and n = 5 mice per group. (f) Schematic showing timeline of administration of anti-NK1.1 antibody and CAR-T cells for *in vivo* NK cell depletion in the presence of CAR-T therapy. (g) Addition of NK1.1 antibody significantly extends survival in mCD19-CAR-T treated mice compared to all other cohorts. Significance for survival experiments was determined using log-rank tests. For both (e) and (g) n = 5-8 mice per group. *P<0.05; **P < 0.01; ***P < 0.001; ****P < 0.001.

**Supplementary Figure 7.**
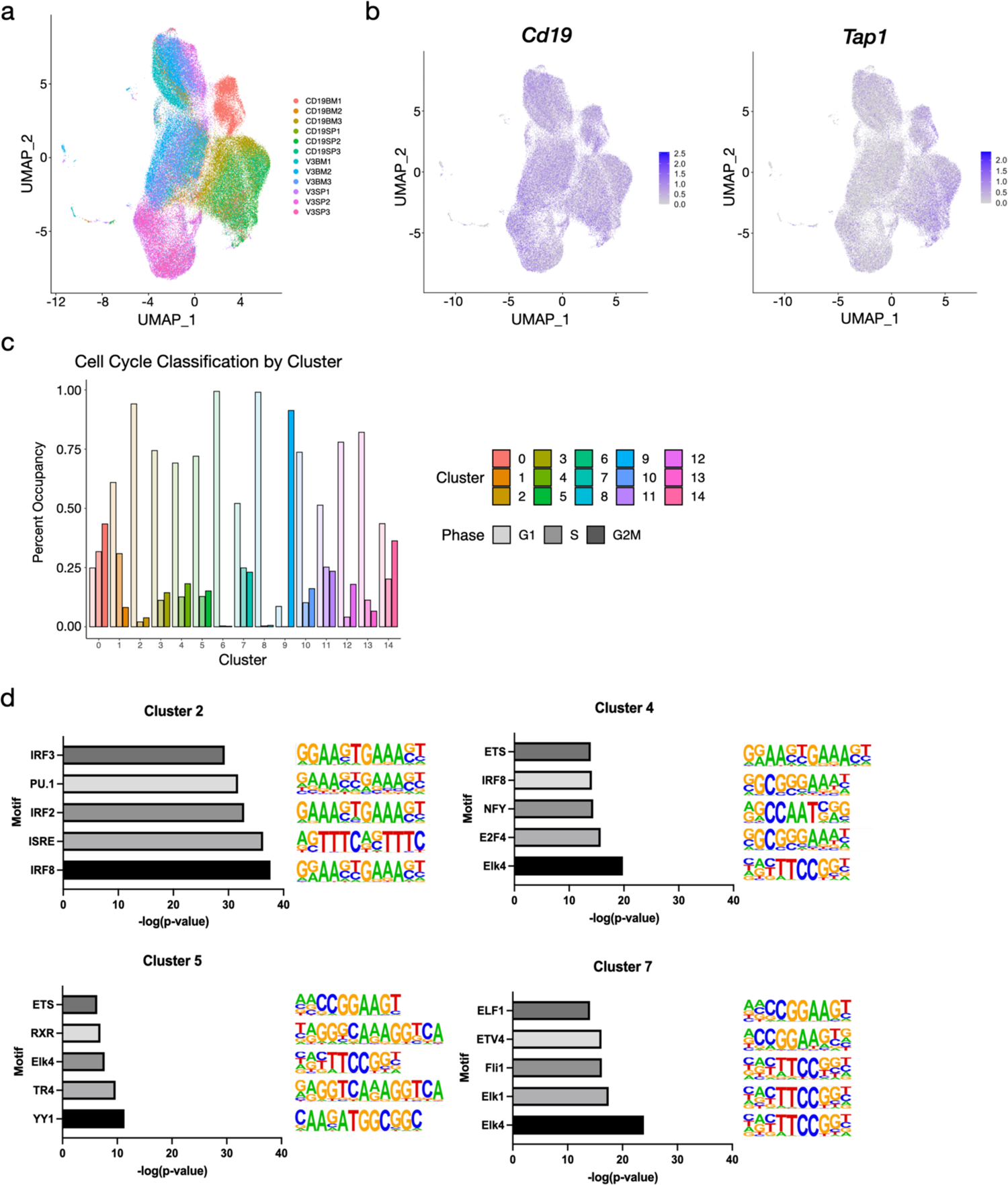
Additional characterizations of clusters that emerge from single-cell expression profiling of CAR-T cell treated animals. (a) 2-dimensional UMAP plots of single cell gene expression profiles collected from mice treated with either anti-mCD19 or anti-hEGFRvIII CAR T cell therapy, with cells color-coded by each individual mouse in the dataset. (b) UMAP plot showing expression of *Cd19* (left panel) and *Tap1* (right panel) in the single cell expression dataset. (c) Proportions of cells assigned to each cell cycle stage in each of the 15 clusters identified in the single cell expression data. (d) Transcription factor motif enrichment analysis for signature genes in clusters 2, 4, 5 and 7. Shown are -log10 transformed *p*-values for top 5 enriched motifs along with their position-weight matrix logos for each cluster.

**Supplementary Figure 8.**
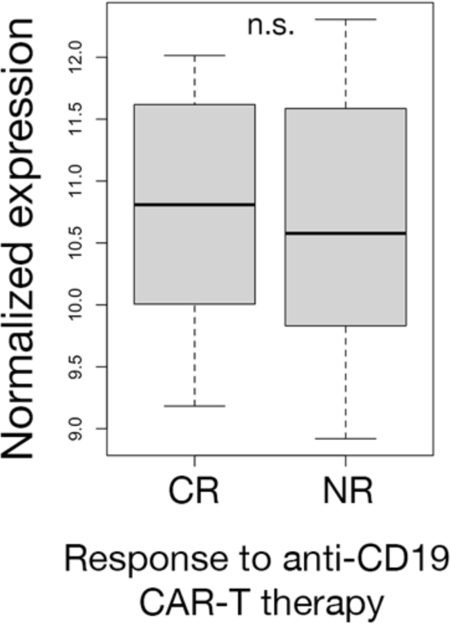
Expression of HLA-E in pre-treatment samples of patients with B-ALL does not correlate with clinical outcome. Normalized expression of HLA-E in pre-treatment patient samples from complete responders (CR) and non-responders (NR) do not significantly differ. Significance is determined using unpaired two-sided student’s t-test. Data are mean ± s.e.m. *P<0.05; **P < 0.01; ***P < 0.001; ****P < 0.001.

