## Supplemental full wester blot figures for "Leukemia-intrinsic determinants of CAR-T response revealed by iterative *in vivo* genome-wide CRISPR screening"

### Supplementary Figure 1a
### Full Western Blots

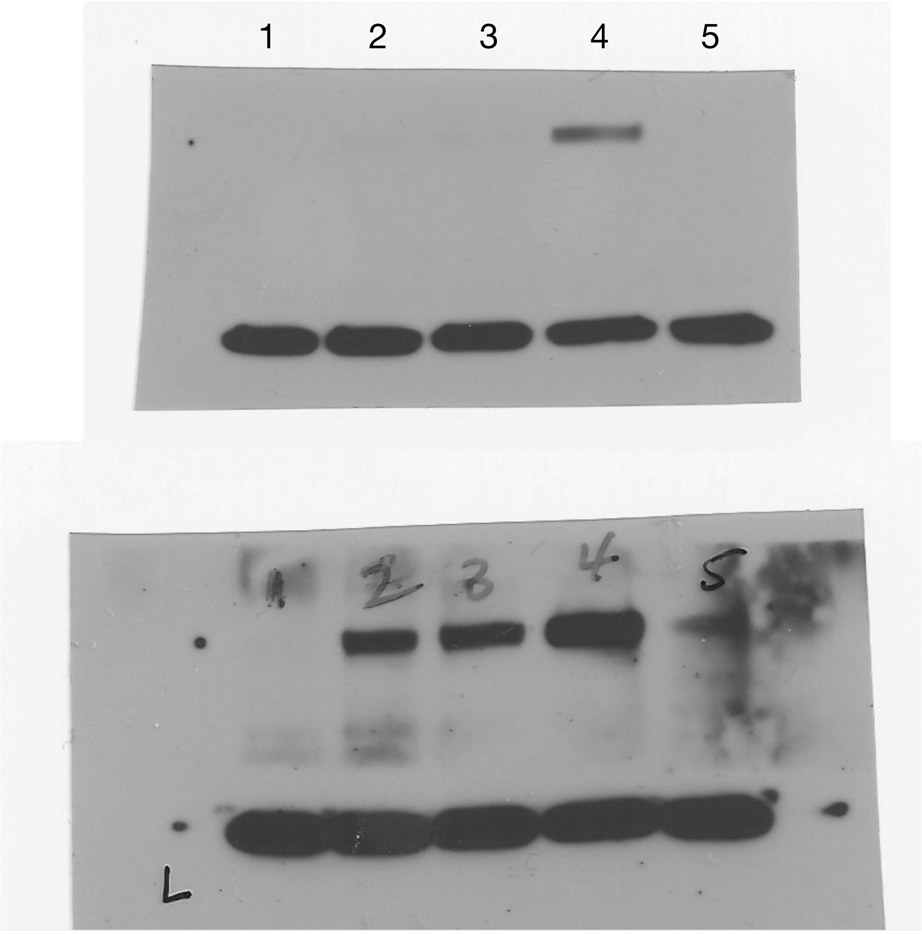


Low exposure (20 sec)

High exposure (60 sec)

1. B-ALL Tomato$^+$ Cas9$^-$
2. Clone 20.8 Tomato$^+$ Cas9$^+$
3. Clone 20.8 Tomato$^+$ GFP$^+$ Cas9$^+$
4. Clone 20.12 Tomato$^+$ GFP$^+$ Cas9$^+$
5. B-ALL Tomato$^+$ GFP$^+$ Cas9$^+$ (Bulk infected)

Supplementary Figure 3b
Full Western Blots

1 2 3 4 5 6

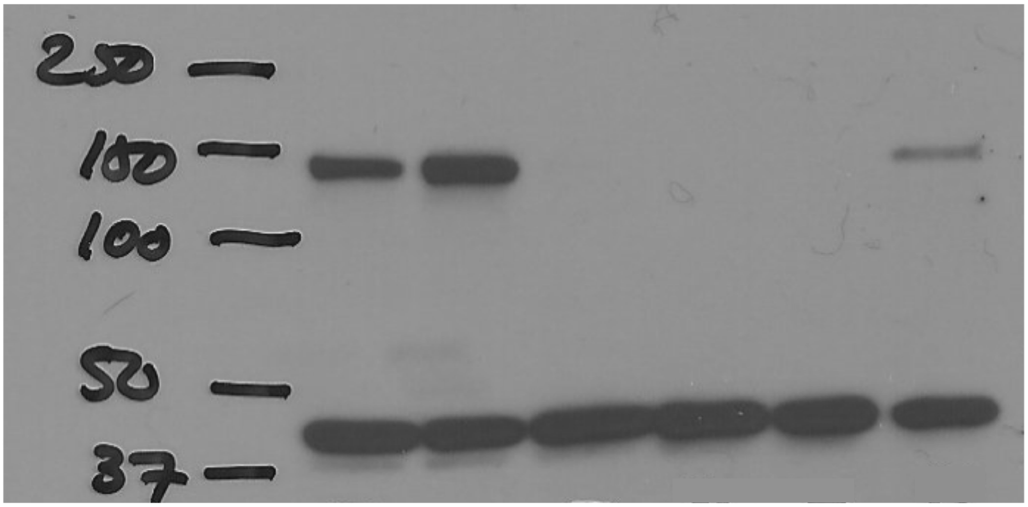


Low exposure (1 min)

1 2 3 4 5 6

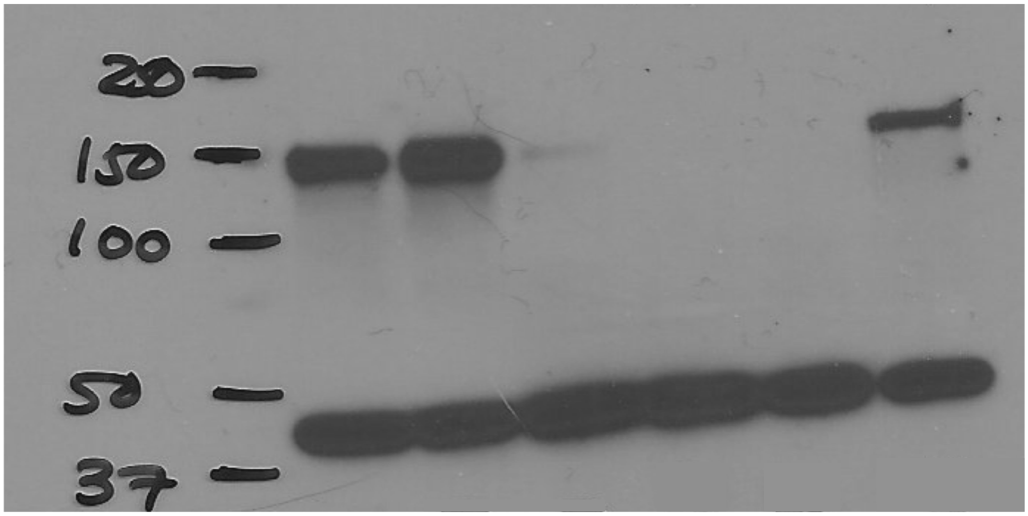


High exposure (2 min)

1. Clone RH53 Tomato$^{+}$ Cas9$^{+}$
2. Clone 20.12 Tomato$^{+}$ GFP$^{+}$ Cas9$^{+}$
3. Clone RH64 Tomato$^{+}$ Cas9$^{Low}$
4. B-ALL Tomato$^{+}$ Cas9$^{-}$
5. B-ALL Tomato$^{+}$ GFP$^{+}$ Cas9$^{-}$
6. Clone RH62 Tomato$^{+}$ Cas9$^{+}$

### Supplementary Figure 3d
#### Full Western Blots

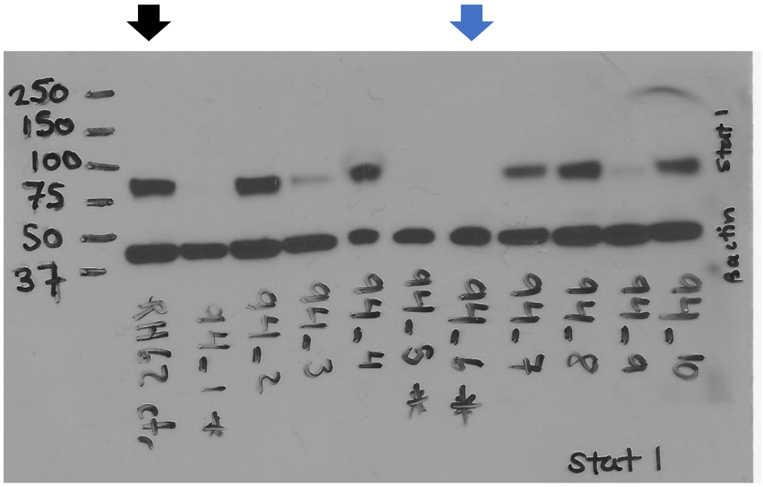


STAT1 (87 kDa)

β-Actin (45 kDa)

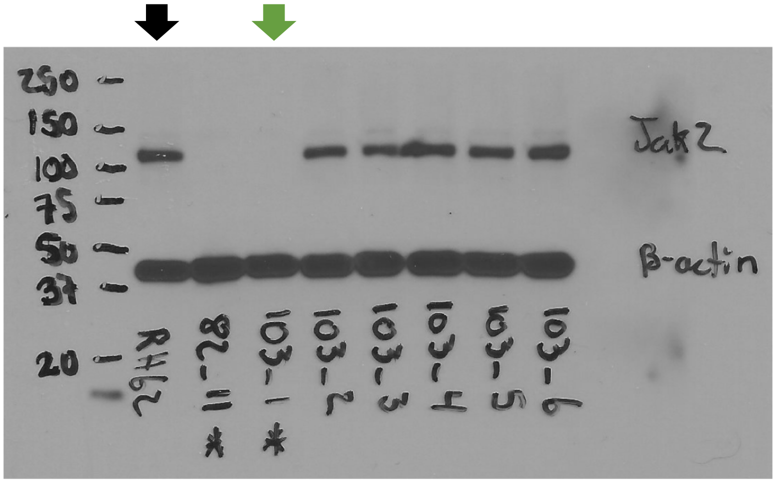


JAK2 (125 kDa)

β-Actin (45 kDa)

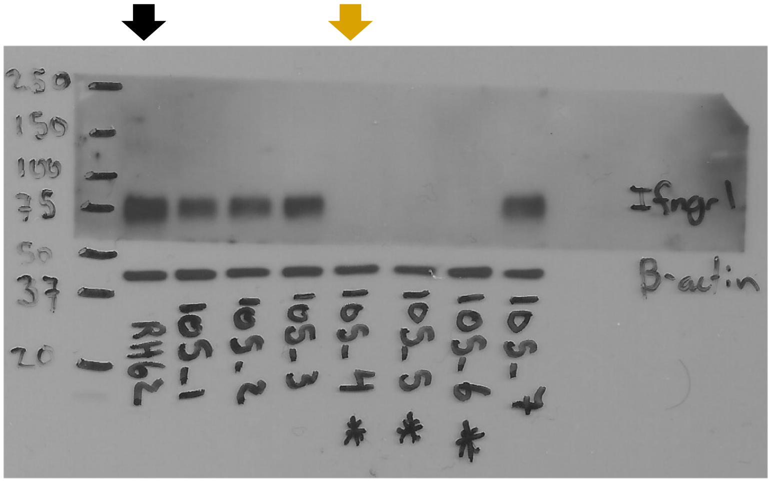


IFNGR1 (70-95 kDa)

β-Actin (45 kDa)

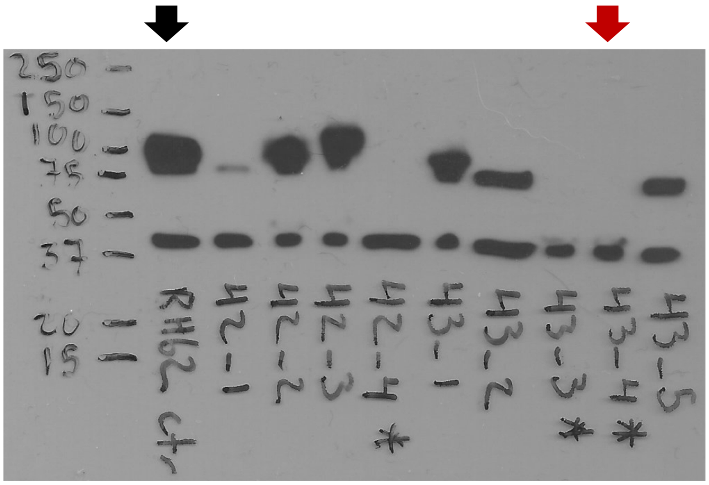


CD19 (95 kDa)

β-Actin (45 kDa)

- WT CTR scc#1
- *Stat1*$^{-/-}$ scc#2
- *Jak2*$^{-/-}$ scc#3
- *Ifngr1*$^{-/-}$ scc#4
- *Cd19*$^{-/-}$ scc#5
